# Environmental Optimization Enables Maintenance of Quiescent Hematopoietic Stem Cells Ex Vivo

**DOI:** 10.1101/475905

**Authors:** Hiroshi Kobayashi, Takayuki Morikawa, Ayumi Okinaga, Fumie Hamano, Tomomi Hashidate-Yoshida, Shintaro Watanuki, Daisuke Hishikawa, Hideo Shindou, Fumio Arai, Yasuaki Kabe, Makoto Suematsu, Takao Shimizu, Keiyo Takubo

## Abstract

Hematopoietic stem cells (HSCs) maintain lifelong hematopoiesis by remaining quiescent in the bone marrow niche. Recapitulation of a quiescent state in culture has not been achieved, as cells rapidly proliferate and differentiate in vitro. After exhaustive analysis of different environmental factor combinations and concentrations as a way to mimic physiological conditions, we were able to maintain engraftable quiescent HSCs for 1 month in culture under very low cytokine concentrations, hypoxia, and very high fatty acid levels. Exogenous fatty acids were required likely due to suppression of intrinsic fatty acid synthesis by hypoxia and low cytokine conditions. By contrast, high cytokine concentrations or normoxia induced HSC proliferation and differentiation. Our novel culture system provides a means to evaluate properties of steady state HSCs and test effects of defined factors *in vitro* under near-physiological conditions.

## INTRODUCTION

Hematopoietic stem cells (HSCs) maintain lifelong hematopoiesis at steady state and after stress by producing multipotent progenitors (MPPs), lineage committed progenitors, and their differentiated progeny within bone marrow niche (Morrison and Scadden, 2014). Cell cycle quiescence distinguishes HSCs from other hematopoietic progenitors and protects them from stress encountered following bone marrow transplant, chemotherapy and radiation (Trumpp et al., 2010). HSC capacity can be assessed by combining transplantation assays with prospective isolation by fluorescence-activated cell sorting (FACS). Recent studies, however, indicate that when unperturbed HSCs behave differently than they do in a transplantation setting (Busch et al., 2015; Sun et al., 2014), possibly due to hyperactivation of cytokine signaling, changes in oxygen status, or lack of niche components.

HSC culture *in vitro* is the subject of intense research due to the utility of HSCs to clinical and basic science applications (Pineault and Abu-Khader, 2015). Functional HSCs have been successfully maintained or expanded in multiple conditions, including treatment with specific cytokines or signaling factors (Delaney et al., 2010; Ema et al., 2000; Gammaitoni et al., 2003; Rollini et al., 2004; Rossmanith et al., 2001), culture in a hypoxic environment (Danet et al., 2003; Hammoud et al., 2012; Mantel et al., 2015; Shima et al., 2009), co-culture with niche cells (Butler et al., 2012; Isern et al., 2013; Yamaguchi et al., 2001), transduction with critical regulators (Antonchuk et al., 2002; Kyba et al., 2002), treatment with SIRT1-inhibitory small molecules, copper chelation, inhibition of aryl hydrocarbon receptors, and activation of glycolysis (Boitano et al., 2010; Fares et al., 2014; Guo et al., 2018; Peled et al., 2004; Peled et al., 2012; Wagner et al., 2016). Some of these methods have been tested in clinical trials and shown to promote more durable engraftment (Pineault and Abu-Khader, 2015). However, although we know that HSCs differ from proliferating HSCs in terms of cell cycling, metabolism, differentiation potential, and stress resistance (Chandel et al., 2016; Ito and Suda, 2014; Trumpp et al., 2010), less is known about how HSCs can be kept quiescent *in vitro*.

Hypothesizing that conditions favoring quiescence are present in bone marrow but lacking in many culture systems, we aimed to define a minimal set of factors that would mimic the bone marrow microenvironment and keep HSCs quiescent and functional in vitro. We show here that HSCs have a unique nutrient requirement for fatty acids. We also optimized environmental conditions, including low cytokine concentrations and maintenance of hypoxia, to favor maintenance of undifferentiated and quiescent HSCs. These conditions were equally applicable to both murine and human HSCs. Development of this system opens an avenue to study steady state HSC properties and manipulate defined factors *in vitro*.

## RESULTS

### Fatty acid biosynthesis is insufficient in HSCs cultured in serum-free culture conditions

To assess potential changes in gene expression in HSCs after conventional culture, we performed cDNA microarray analysis comparing HSCs immediately after sorting and HSCs cultured for 16 hours with SCF and TPO with or without serum (Figure 1A). Genes most significantly upregulated in serum-free culture conditions were fatty acid and cholesterol biosynthesis-related genes (Figures 1B-1D and S1A), a finding validated by qPCR (Figure 1F). We also observed upregulation of cell division-related genes and downregulation of stem cell-related genes in cultured versus fresh HSCs, irrespective of the presence or absence of serum (Figure S1B). Relevant to lipid metabolism, target genes of the master regulator sterol regulatory element-binding protein SREBP (Figure 1E) were significantly enriched in HSCs cultured in serum-free conditions compared with freshly sorted HSCs or HSCs cultured in 10% serum (Figures 1C, S1C, and S1D). Three SREBP isoforms activate partially overlapping targets: SREBP-1c activates primarily fatty acid synthesis genes, SREBP-2 activates cholesterol-related genes, and SREBP-1a activates both (Eberlé et al., 2004). Our findings suggest that upregulation of SREBP targets in serum-free medium may be triggered by a relative shortage of fatty acids or cholesterol. Thus we initially focused on effects of fatty acids on HSC proliferation. To do so, we cultured HSCs in either serum-free or 10% serum conditions and blocked various steps of fatty acid biosynthesis using the inhibitors TOFA, cerulenin, or A939572. HSC proliferation in serum-free conditions was decreased by inhibition of any step in fatty acid synthesis, whereas addition of 10% serum rescued proliferation activity in the presence of any of these inhibitors (Figure 1G). These results show that HSCs require fatty acids for proliferation, either through intrinsic biosynthesis or from an exogenous source.

**Figure 1.**
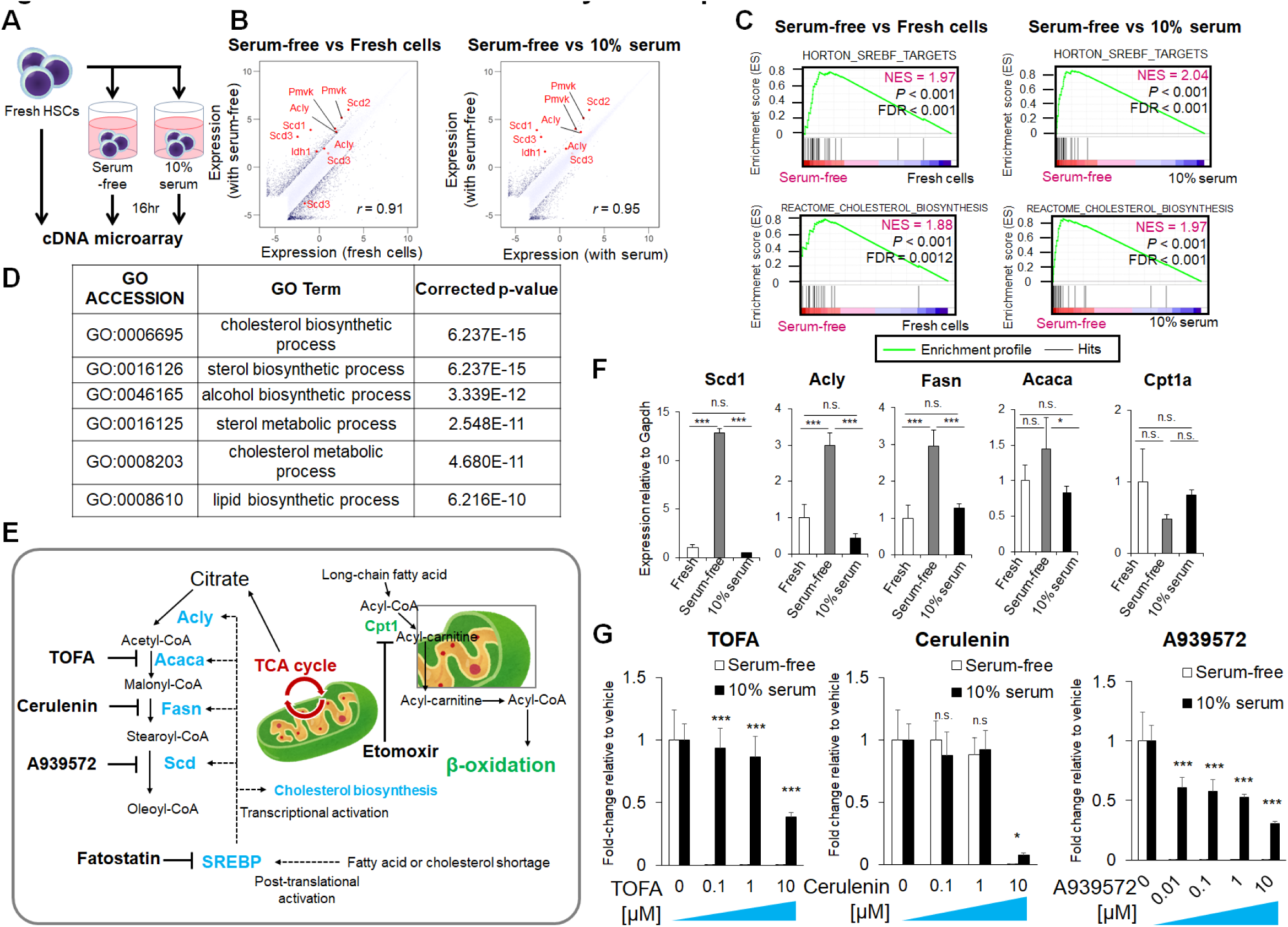
Serum-free culture induces fatty-acid dependence in HSCs. (**A**) Schematic showing microarray analysis. Freshly isolated CD150^+^CD48^-^CD41^-^Flt3^-^CD34^-^ LSK cells (Fresh HSCs), and HSCs cultured 16 hours with (10% serum) or without (Serum-free) serum were subjected to cDNA microarray analysis. Cytokines (SCF and TPO at 100ng/mL each) were supplemented in each condition. n =1 for each condition. (**B**) Scatter plot of cDNA microarray data comparing conditions indicated in (**A**). Fatty acid or cholesterol synthesis genes are highlighted in red. Correlation coefficient *r* is noted. (**C)** Gene set enrichment analysis (GSEA) of cDNA microarray data of HSCs shown in (**A**). FDR, false discovery rate. NES, normalized enrichment score. (**D**) Gene ontology analysis (GO) of HSCs indicated in (**A**). Shown are GO terms of genes over-represented in HSCs cultured in serum-free conditions. (**E**) Schematic illustration of fatty acid biosynthetic and β-oxidation pathways, and their small molecule inhibitors used here. Regulators related to fatty-acid biosynthesis are in blue and to β-oxidation are in green. Arrows indicate enzymatic reactions and dotted arrows indicate transcriptional and post-translational activation. (**F**) Quantitative real-time PCR (qPCR) analysis of fatty acid-related mRNAs in fresh HSCs and HSCs cultured 16 hours with or without 10% serum. Values are normalized to Gapdh expression and expressed as fold-induction relative to levels in fresh HSCs (mean ± SD, n = 4 technical replicates). \**P* < 0.05, \*\**P* < 0.01, \*\*\**P* < 0.001, by Tukey-Kramer multiple comparison test. (**G**) Relative survival of cultured HSCs following treatment with indicated inhibitors. 100 HSCs were cultured 7 days in serum-free or 10% serum conditions. Cytokines (SCF and TPO at 100ng/mL each) were added to each culture. Data are normalized to vehicle control (mean ± SD, n = 4). \**P* < 0.05 and \*\*\**P* < 0.001 by two-tailed Student’s t test between Serum-free sample and serum conditions.

### Albumin serves as a fatty acid source *in vitro*

Most circulating fatty acid is contained in lipoproteins, which are partially hydrolyzed to produce free fatty acids, which are then bound to albumin as a carrier (Spector, 1975). To determine whether circulating albumin diffuses into the bone marrow cavity as a fatty acid source *in vivo*, we used intravital imaging and multi-photon microscopy to determine whether FITC-labeled bovine-serum albumin (BSA; MW 66.5kDa) penetrates the extravascular space of cranial bone marrow. When we injected mice with labeled BSA, we observed that it gradually diffused into the bone marrow extravascular space over 60 minutes of observation (Figures 2A and 2B). By contrast, that diffusion pattern was not observed following comparable injection 150kDa dextran-FITC (Figures 2A and 2B), suggesting that bone marrow vasculature acts as a size barrier but allows diffusion of circulating albumin, in line with previous reports (Michelsen, 1969; WASSERMAN and MAYERSON, 1951). Using ELISA analysis, we found that albumin concentration in bone marrow extracellular fluid was 13 mg/mL, approximately one third that of serum albumin concentration (Figure S2A). Thus, we chose to utilize BSA as a fatty acid source in *in vitro* culture. As a first step, we used gas chromatography-mass spectrometry to quantify fatty acid bound to commercially available batches of BSA (hereafter designated native BSA (nBSA)) by a lipidomics approach. We observed that saturated and unsaturated fatty acids bound to various lots of nBSA, while levels of fatty acid bound to BSA defined as “fatty acid-free” (FA-free) were almost negligible (Figure S2B). We then cultured HSCs and MPPs *in vitro* to evaluate effects of nBSA added to serum-free SF-O3 medium. Addition of nBSA to the medium promoted HSC and MPP proliferation dose-dependently and to a greater extent than culture in 10% serum (Figure S2C). We also observed decreased expression of SREBP-target genes after 16 hours of culture with 4% nBSA, with reconstituted BSA (defined as BSA reconstituted with sodium palmitate and sodium oleate), or with 10% serum relative to expression levels in 0.1% nBSA or FA-free BSA conditions both in HSCs (Figure 2C) and MPPs (Figure S2E). *Srebf*-*1* and *-2* mRNA levels were suppressed by 10% serum, indicating that SREBP activity is partially regulated at the transcriptional level in HSCs (Figure S2D). Notably, addition of 4% nBSA or reconstituted BSA suppressed expression of not only fatty acid synthesis genes but of Pmvk, a cholesterol synthesis gene, suggesting that fatty acid deficiency may upregulate expression of cholesterol-related genes (Figures 2C and S2E).

**Figure 2.**
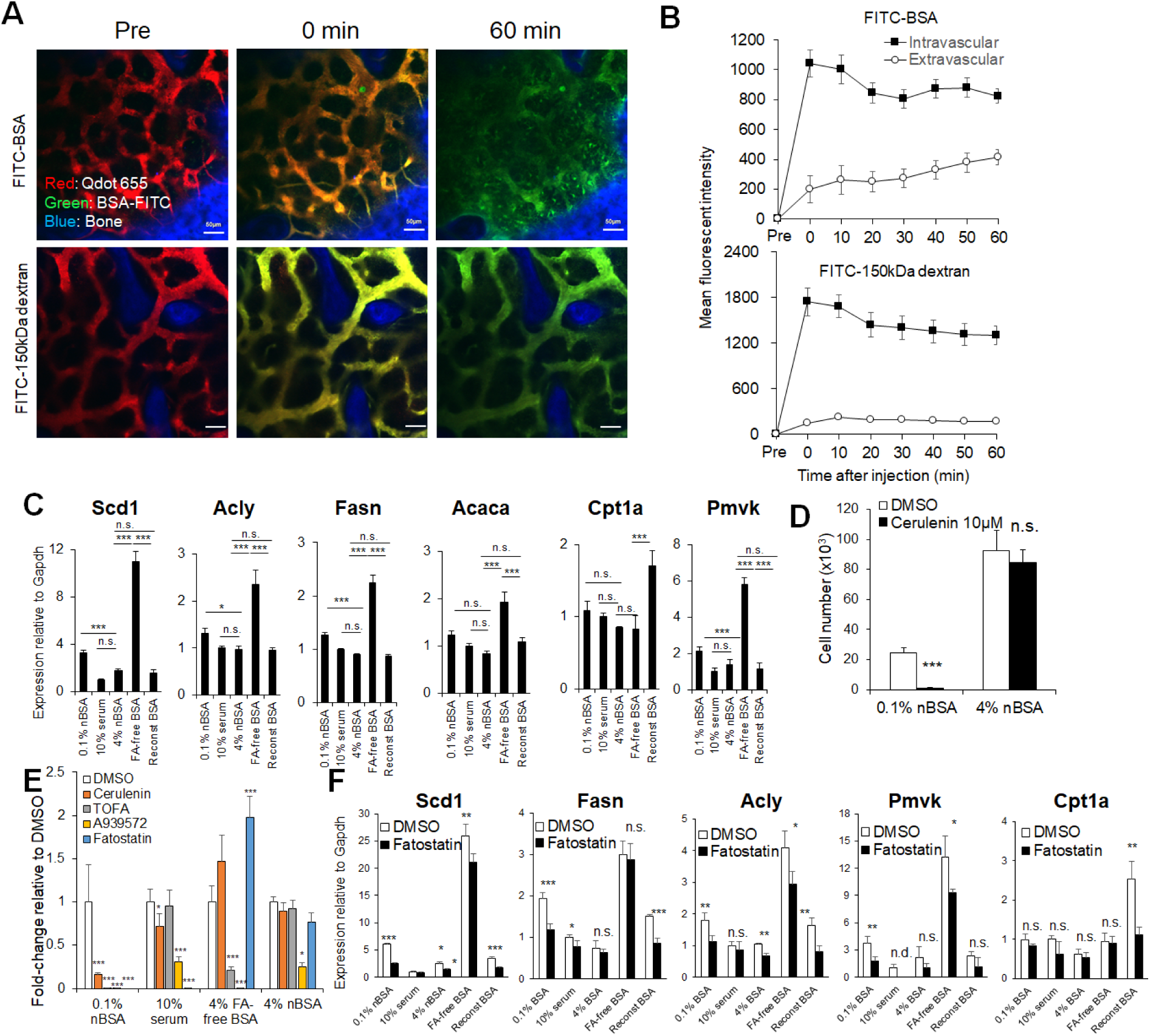
Albumin is a source of fatty acid for HSCs. (**A**) Representative intravital staining imaging of mouse calvarial bone marrow using multi-photon excitation microscopy to assess albumin permeability. Qdot-655 (red) was administered immediately prior to intravenous injection of FITC-labelled BSA (green, top panels) or FITC-labelled 150kDa dextran (green, bottom panels) to visualize vasculature. Images were acquired at indicated representative time points after BSA or dextran injection (Pre, prior to BSA or dextran injection). Blue indicates second harmonic generation overlapping with bone area. (**B**) Mean fluorescent intensity (MFI) of the FITC-BSA (top) or FITC-150kDa dextran (bottom) signal in intra- and extra-vascular regions of calvarial bone marrow, based on microscopy analysis shown in (**A**) (mean ± SD of 18 randomly chosen points for each indicated area from 3 mice from 3 independent experiments). (**C**) qPCR analysis of fatty acid- or cholesterol-related mRNAs in CD150^+^CD41^-^CD48^-^Flt3^-^CD34^-^ LSK cells cultured 16 hours with indicated reagents plus cytokines (SCF and TPO at 100ng/mL each). nBSA, native bovine serum albumin. FA-free BSA: 4% w/v fatty acid-free BSA. Reconst BSA: 4% w/v fatty acid-free BSA reconstituted with sodium palmitate and sodium oleate at 200μg/mL. Values are normalized to Gapdh expression and expressed as fold-induction relative to levels in 10% serum (mean ± SD, n = 4 technical replicates). \**P* < 0.05, \*\**P* < 0.01, \*\*\**P* < 0.001. n.s., not significant by Tukey-Kramer multiple comparison test. (**D**) Total cell number after 7 days of culture of 100 CD150^+^CD41^-^CD48^-^Flt3^-^CD34^-^ LSK cells with or without 10μM cerulenin (mean ± SD, n = 4). Media contained 0.1% or 4% (w/v) nBSA plus SCF and TPO at 100ng/mL each. \*\*\**P* < 0.001. n.s., not significant by two-tailed Student’s t test between DMSO control and Cerulenin. (**E**) Relative total cell number after 7 days of culture of 100 CD150^+^CD41^-^CD48^-^Flt3^-^CD34^-^ LSK cells with indicated fatty acid biosynthesis inhibitors under different culture conditions (mean ± SD, n = 4 from 1 experiment). Cerulenin, 5μM; TOFA 1μM; A939572, 1μM; and Fatostatin, 20μM. Final DMSO concentration, 0.1% v/v. Data are normalized to cell numbers seen in DMSO controls. \**P* < 0.05, \*\**P* < 0.01, and \*\*\**P* < 0.001. n.s., not significant compared with DMSO by Tukey-Kramer multiple comparison test. (**F**) qPCR analysis of fatty acid or cholesterol-related mRNAs in CD150^+^CD41^-^CD48^-^Flt3^-^CD34^-^ LSK cells cultured 16 hours with or without 20μM Fatostatin under indicated conditions. Indicated cultures were supplemented with SCF and TPO at 100ng/mL each. DMSO concentration, 0.1% v/v. nBSA, FA-free BSA, and Reconst BSA are defined as in **(C).** Values were normalized to Gapdh expression and expressed as fold-induction relative to levels detected seen in DMSO/10% serum sample (mean ± SD, n = 4 technical replicates from 1 experiment). \**P* < 0.05, \*\**P* < 0.01, and \*\*\**P* < 0.001. n.s., not significant by two-tailed Student’s t test, n.d. not detected.

Addition of 4% nBSA to mimic serum albumin levels restored HSC proliferation capacity in the presence of inhibitors of fatty acid synthesis to a level similar to 10% serum conditions, as compared to 0.1% nBSA or 4% FA-free BSA (Figures 2D, and 2E). Effects of fatty acid inhibitors differed with culture conditions. For example, fatostatin, which blocks post-translational SREBP activation, strongly suppressed proliferation of HSC-derived colonies in the 10% serum group but not in the FA-free BSA group. Treatment with cerulenin did not alter HSC-derived colony growth in the FA-free BSA group compared with 0.1% nBSA group. SREBP target genes were more highly upregulated in the FA-free BSA group, and fatostatin effects on this upregulation were minimal (Figure 2F). In contrast, fatostatin effectively suppressed expression of SREBP target genes in the 0.1% nBSA group (Figure 2F). Overall, we conclude that BSA serves as a fatty acid source in HSC culture and can replace serum in terms of fatty acid supply. Hereafter we utilized 4% nBSA in HSC culture medium.

### Lower cytokine levels and hypoxia maintain phenotypic HSCs

Cytokine combinations that favor HSC quiescence *in vitro* remain unclear, although it is known that hypoxia is also critical for quiescence (Cipolleschi et al., 1993; Takubo et al., 2010), with the average physiological oxygen concentration as low as 1.3% around sinusoid (Spencer et al., 2014). Thus, to determine optimal culture conditions in the presence of essential cytokines we assessed differentiation and proliferation following 7 days of culture in various concentrations of SCF (0.33-81ng/mL, or none) and TPO (0.02-62.5ng/mL, or none) plus 4% BSA, in hypoxic (1% O_2_) or normoxic (20% O_2_) conditions. As expected, we observed that total numbers of cells derived from HSCs increased in parallel with SCF and TPO levels, indicating that SCF and TPO promote cell cycling (Figures 3A and S3A). Moreover, at any cytokine concentration, total cell numbers were lower in hypoxia, as reported (Cipolleschi et al., 1993). Under hypoxia, the number of megakaryocytes was highest at high TPO/low SCF conditions, while under normoxia, the number of megakaryocytes increased with increasing SCF and TPO levels (Figures 3B and S3B). Notably, under either hypoxia or normoxia, cytokine concentrations were negatively correlated with phenotypic HSC frequency, defined as the CD150^+^CD48^-^ fraction in Lineage marker^-^Sca1^+^c-Kit^+^ (LSKs), suggesting that HSCs proliferated and differentiated under high cytokine and normoxic conditions (Figures 3C and S3C). LSK fractions were divided into 4 subgroups based on CD150 and CD48 expression and the number of cells in each fraction was determined (Figures 3D and S3D). Higher concentrations (>27g/mL SCF) promoted HSC loss (Figures 3D and S3D). On the other hand, high TPO (62.5ng/mL) and low SCF (<3ng/mL) conditions most favored an increase in the number of phenotypic HSCs, with approximately 40% of cells remaining within the CD150^+^CD48^-^ LSK fraction and increasing 2-fold in number. The number of CD150^-^CD48^-^ LSK cells (phenotypic MPP1s) increased with SCF levels, but was not altered by changes in TPO levels, whereas the number of CD150^+^CD48^+^ LSK cells (phenotypic MPP2s) was increased only by increasing TPO concentration (Figures 3D, S3D, and S3E). We conclude that in vitro the first bifurcation of HSC differentiation into CD150-negative or CD48-positive fractions is determined by differential responses to SCF or TPO, respectively. Increased partial oxygen pressure also increased the number of the differentiated CD150^+^CD48^+^ and CD150^-^CD48^+^ LSK cells. These patterns differed significantly under 0.1% BSA conditions: in that case, HSC numbers were highest at 100ng/mL SCF and 100ng/mL TPO, but cells survived at lower (< 3ng/mL SCF) cytokine conditions (Figure S3F).

**Figure 3.**
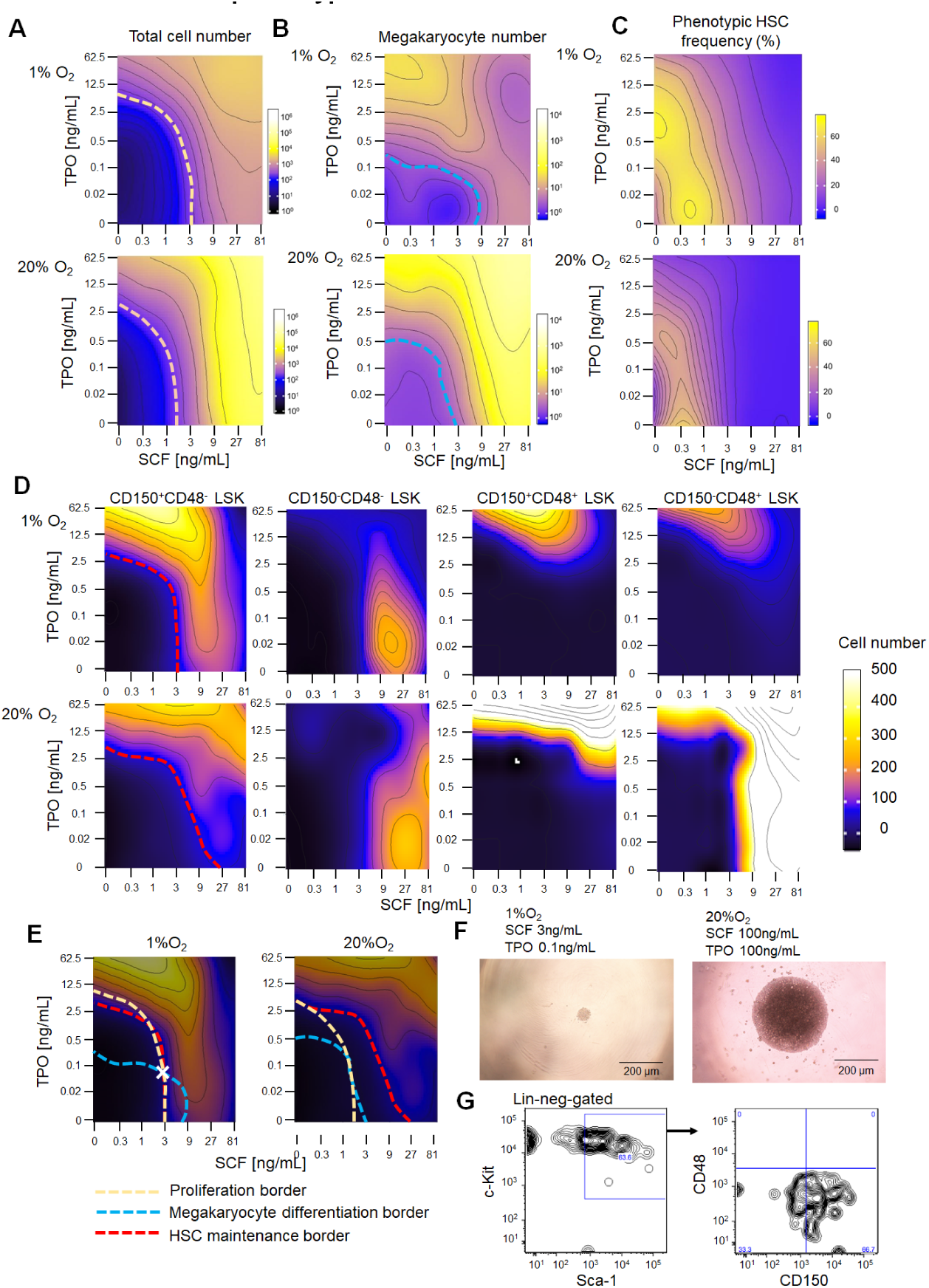
Hypoxia and low cytokine conditions are essential for maintenance of phenotypic HSCs. (**A**-**D**) Contour plots showing cell number or frequency after 7 days culture (input: 275-300 CD150^+^CD41^-^ CD48^-^Flt3^-^CD34^-^ LSK cells) in 1% or 20% O2 conditions in 49 (7 x 7) different concentrations of SCF (0 to 81 ng/mL) and/or TPO (0 to 62.5ng/mL). Total cell number (**A**), megakaryocyte (forward scatter^hi^, CD150^hi^ and CD41^hi^) number (**B**), phenotypic HSC (CD150^+^CD48^-^ LSK cells) frequency per total cells (**C**), and absolute number of each LSK subfraction (**D**) are shown (n = 3 for each group from 3 independent experiments). In (**A**), yellow dashed line demarcates the point at which cells remain quiescent (cell number ∼300). In (**B**), blue dashed line demarcates that point at which megakaryocyte number is ∼10. In (**D**), red dashed line demarcates the point at which the number of phenotypic HSCs is maintained (∼100 cells). (**E**) Optimization of cytokine and oxygen conditions for HSC maintenance. Proliferation, megakaryocyte differentiation, and HSC maintenance borders (as indicated in (**A**), (**B**), and (**D**), respectively) are merged, and intersection those borders observed in 1% O_2_ is marked by “X”. Note that proliferation and HSC maintenance borders differ in 20% O_2_ conditions. (**F**) Representative images of HSCs cultured 7 days in either 1% O_2_ with 3ng/mL SCF and 0.1ng/mL TPO, or with 20% O_2_ with SCF and TPO at 100ng/mL each. Bars, 200μm. (**G**) Representative plot of flow cytometry of HSCs cultured 7 days in 1% O_2_ with 3ng/mL SCF and 0.1ng/mL TPO.

In vivo HSCs remain generally quiescent with minimal output of differentiated cells. Thus, we assessed HSC maintenance *in vitro* based on 3 criteria: 1) comparison of total cell number at the start and after 7 days of culture (starting with 300 cells), 2) a minimal (∼10 cells) megakaryocyte number, and 3) the presence of ∼100 phenotypic HSCs. Those conditions were met at 3ng/mL SCF, 0.1ng/mL TPO, and 1% O_2_ conditions, while at 20% O_2_ cells did not maintain quiescence even at low cytokine conditions (Figure 3E). Colonies of cultured HSCs under optimal conditions appeared much smaller than those grown in 100ng/mL SCF, 100ng/mL TPO, and 20% O_2_ conditions (Figure 3F). Greater than 60% of cells in the LSK fraction resided within the CD150^+^CD48^-^ fraction in low cytokine conditions, reflecting minimal differentiation (Figure 3G).

### Near-physiological culture conditions allow maintenance of functional HSCs

Phenotypic HSCs defined by surface markers are not necessarily functional. Thus we characterized function of HSCs cultured in 3ng/mL SCF, 0.1ng/mL TPO, 4% BSA and 1% O_2_ conditions (hereafter, termed “maintenance conditions”, and see Methods for details). We first assessed cell cycle status in maintenance conditions versus “high cytokine” (100ng/mL SCF and 100ng/mL TPO) conditions favoring proliferation under either hypoxia or normoxia. After 16 hours of culture and a 2hr EdU pulse, only 4% of cultured HSCs under maintenance conditions incorporated EdU in contrast to 40% under proliferation conditions (Figures 4A and 4B). Oxygen concentration slightly activated cell cycle in this time frame, although not significantly (Figures 4A and 4B). The frequency of apoptotic Annexin V^+^ propidium iodide (PI)^-^ cells was 5% both in 4% nBSA and in the reconstituted BSA groups, and 15% in FA-free BSA group, confirming that fatty acids are required for cell survival in maintenance conditions (Figure 4C).

**Figure 4.**
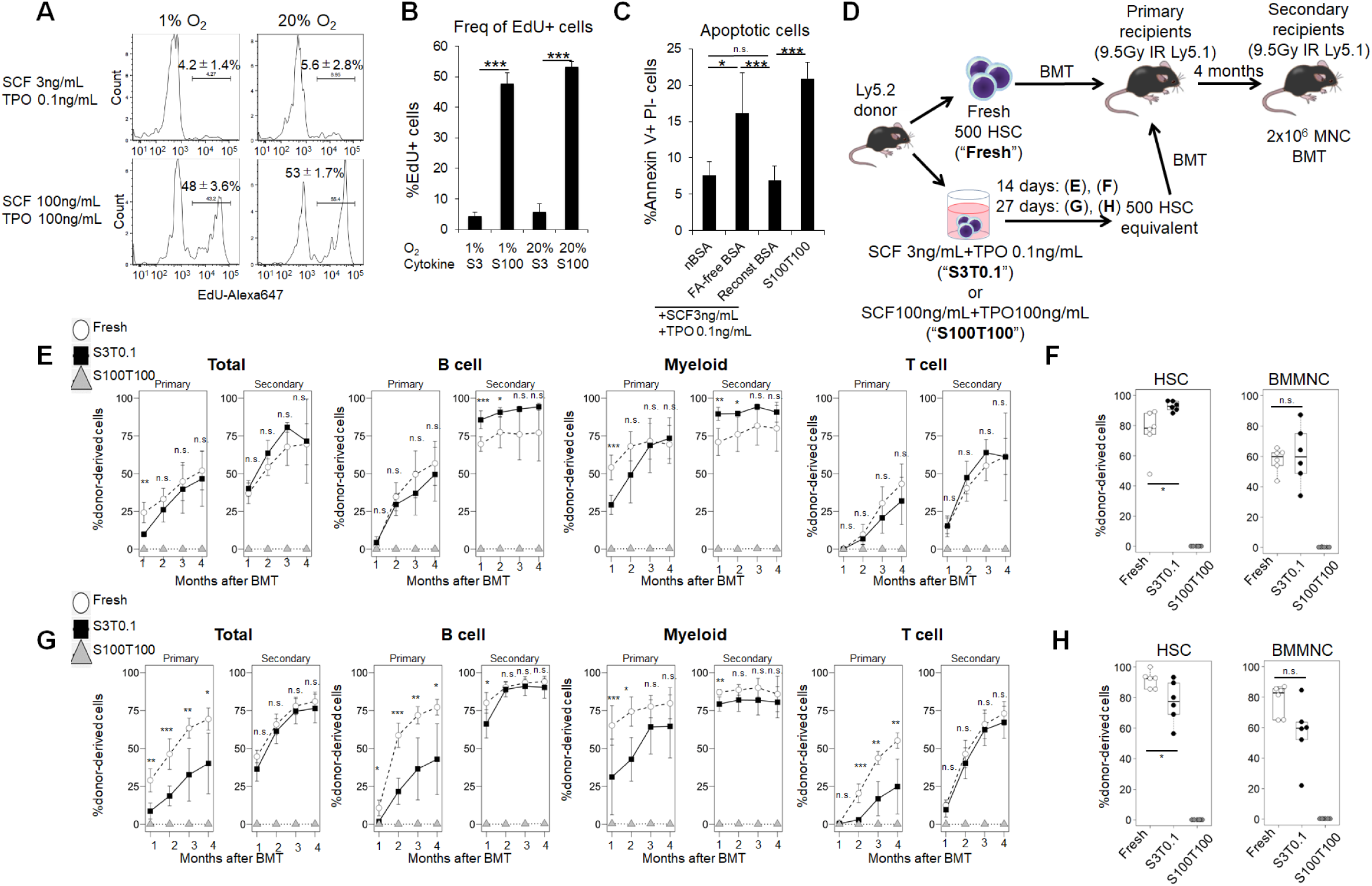
Hypoxia and low cytokine conditions are essential for maintenance of functional HSCs. (**A** and **B**) EdU incorporation in HSCs cultured 16 hours under indicated cytokine and O_2_ conditions. Cells were EdU-labeled for 2 hr (**A**) and analyzed by flow cytometry (mean ± SD, n = 4). Representative histograms show frequency of EdU-incorporated cells (**B**). \**P* < 0.05, \*\**P* < 0.01, and \*\*\**P* < 0.001. n.s., not significant by Tukey-Kramer multiple comparison test. (**C**) Apoptosis assay of HSCs cultured 5-days in indicated conditions. nBSA: 4% w/v native BSA. FA-free BSA: 4% w/v fatty acid-free BSA. Reconst. BSA: 4% w/v fatty acid-free BSA reconstituted with 200μg/mL each of sodium palmitate and sodium oleate (mean ± SD, n = 4). \**P* < 0.05, \*\**P* < 0.01, and \*\*\**P* < 0.001. n.s., not significant by Tukey-Kramer multiple comparison test. (**D-H**) Bone marrow transplant (BMT) of freshly-isolated versus cultured HSCs. HSCs (CD150^+^CD48^-^ CD41^-^Flt3^-^CD34^-^ LSK cells) were sorted and cultured in SF-O3 with 3ng/mL SCF, 0.1ng/mL TPO, in 4% BSA in 1% O_2_ conditions for 14 or 27 days. Then, 500 cultured cells (day 0-equivalent) or 500 freshly-isolated HSCs were transplanted into lethally-irradiated mice, and peripheral blood chimerism was determined. Experimental design (**D**). Peripheral blood chimerism of HSCs cultured 14 days and transplanted into primary or secondary recipients (mean ± SD, n = 6 per each group) (**E**). Bone marrow chimerism of donor-derived cells 4 months after primary BMT in (**E**) (n = 6 per group) (**F**). Peripheral blood chimerism of HSCs cultured 27 days and transplanted into primary or secondary recipients (mean ± SD, n = 6 per each group) (**G**). Bone marrow chimerism of donor-derived cells 4 months after primary BMT in (**G**) (n = 6 per group) (**H**). \**P* < 0.05, \*\**P* < 0.01, and \*\*\**P* < 0.001. n.s., not significant by two-tailed Student’s t test between Fresh and S3T0.1.

To evaluate repopulation capacity, we transplanted fresh HSCs or HSCs cultured 14 or 27 days under maintenance or high cytokine conditions plus 4% nBSA and 1% O_2_ into lethally-irradiated recipient mice (Figure 4D). HSCs cultured in high cytokine conditions could not maintain repopulation capacity when cultured for 14 or 27 days (Figures 4E-4H), as expected from the *in vitro* mapping indicating HSC exhaustion (Figure 3D). HSCs cultured in maintenance conditions for 14 days exhibited behavior almost comparable to that of fresh HSCs during both primary and secondary transplantation (Figure 4E). Donor HSC chimerism within bone marrow 4 months after bone marrow transplantation (BMT) was comparable in fresh and 14-day cultured HSCs (Figure 4F). Following 27-31 days of culture, HSCs showed repopulation capacity at secondary BMT (Figures 4G, S4D and S4E) as well as high bone marrow HSC frequency (Figures 4H and S4F) with chimerism, and lymphoid potential varied slightly among experimental replicates (Figures 4G, and S4D-S4H). Cultured HSCs gave rise to red blood cells (RBCs) and platelets, based on analysis of GFP-expressing donor cells (Figures S4G and S4H). Since the number of phenotypic HSCs increased at high TPO/low SCF conditions (Figure 3D), we also transplanted HSCs cultured with 1ng/mL SCF + 100ng/mL TPO in 1% O_2_ for 11 days. Chimerism of 5 lineages including RBCs and platelets was superior to that of fresh HSCs until 2 months post-transplantation and remained comparable until 4 months (Figures S4A-B, and S4G-H). In contrast, both peripheral blood chimerism at secondary transplantation or bone marrow HSC chimerism of cultured HSCs was less than that of fresh HSCs (Figures S4B and S4C), in agreement with the idea that proliferating HSCs show decreased repopulation capacity (Passegué et al., 2005; Wilson et al., 2008).

To assess fatty acid requirements in maintenance conditions, we cultured HSCs for 28 days in medium containing either 4% nBSA, 4% FA-free BSA, or 4% reconstituted BSA and then performed transplantation. Peripheral blood chimerism in bone marrow was comparable between the nBSA and reconstituted BSA groups, while the FA-free BSA group exhibited no engraftment (Figures S4K and S4L), confirming importance of fatty acids for HSC culture. Notably, although HSC repopulation capacity was maintained in the 3ng/mL SCF and 0.1ng/mL TPO, 1% O_2_, and 4% nBSA condition for 1 month, the number of phenotypic HSCs gradually decreased in maintenance conditions, although it was maintained under 1ng/mL SCF and 100ng/mL TPO conditions (Figure S4M). Also, the frequency of functional HSCs decreased in the maintenance condition as assessed by limiting dilution analysis (data not shown), suggesting that the maintenance condition does not totally recapitulate the *in vivo* microenvironment.

### Multipotent progenitors retain their marker phenotype in maintenance conditions

Multipotent progenitors (MPPs) differ from HSCs in terms of lower self-renewal capacity and higher cell cycling activity (Pietras et al., 2015). If our HSC maintenance conditions faithfully recapitulated the native bone marrow microenvironment, the same conditions should be applicable to the culture of MPPs, as both MPPs and HSCs are maintained in bone marrow. To confirm this, we examined *in vitro* behavior of CD150^-^ CD41^-^CD48^-^Flt3^-^ LSK (MPP1s), CD150^+^CD41/CD48^+^Flt3^-^ LSK (MPP2s), CD150^-^CD41/CD48^+^Flt3^-^ LSK (MPP3s), and Flt3^+^ LSK (MPP4s) grown in HSC maintenance conditions. Surface markers characteristic of MPP1, MPP2, and MPP3 cell types were relatively unchanged after 4 days of culture in SCF 3ng/mL +TPO 0.1ng/mL conditions (Figures S4N and S4O), whereas the number of MPP4 cells greatly decreased, presumably due to lack of essential factors. Addition of as low as 3ng/mL Flt3 ligand (Flt3L) and 0.02ng/mL IL-6 to the maintenance condition, on the other hand, increased MPP4 proliferation to a rate higher than that seen in HSCs under the same culture conditions, a characteristic feature of MPPs (Figures S4P, S4Q, and S4S). By contrast, MPP4 cells were less proliferative than HSCs in medium lacking Flt3L or IL-6, even at higher SCF and TPO concentrations (Figures S4R and S4S), indicating high dependence of MPP4 cells on Flt3L or IL-6.

### Comparison of properties of fresh versus cultured HSCs

As noted, HSCs cultured under maintenance conditions differ from fresh HSCs in terms of lower repopulation capacity following primary transplantation. To understand why, we performed cDNA microarray analysis to capture the transcriptomic profile (Figure 5A) of fresh HSCs versus HSCs cultured under maintenance conditions for 7 days with either nBSA or reconstituted BSA. Global gene expression patterns were similar in fresh HSCs and in HSCs cultured in either nBSA or reconstituted BSA conditions. Upregulation of fatty acid synthesis or cholesterol synthesis genes was, if not perfectly, normalized (Figures 5B, and S5A-B) Gene set enrichment analysis for HSC or cell-cycle genes, which were significantly upregulated in high cytokine condition (Figure S1B), was comparable in fresh and cultured HSCs (Figure 5C). The most prominent difference between fresh and cultured HSCs was upregulation of megakaryocyte-related genes in cultured HSCs under either nBSA or reconstituted BSA conditions. (Figure S5B).

**Figure 5.**
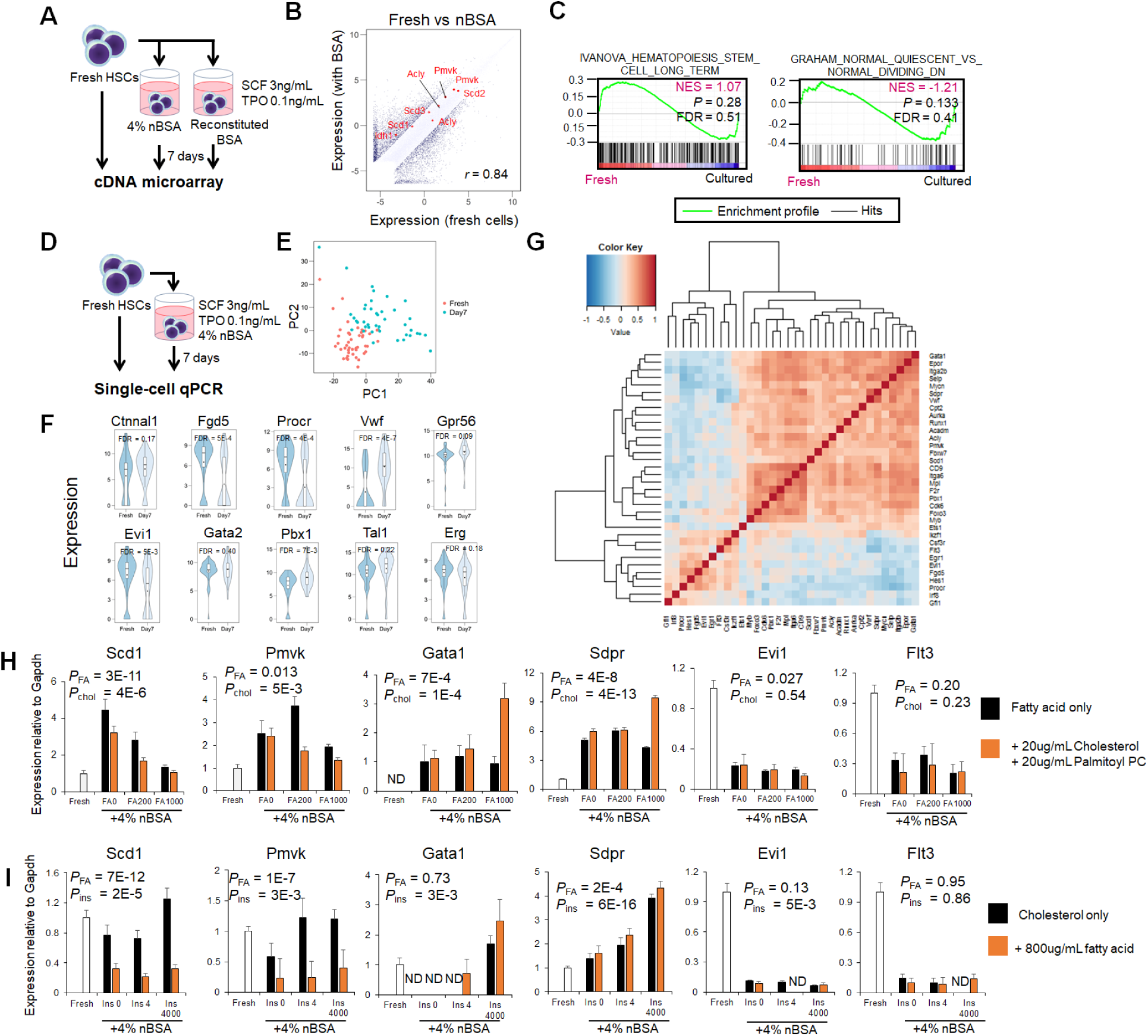
Properties of fresh versus cultured HSCs. (**A**-**C**) cDNA microarray analysis. (**A**) Schematic showing experimental design. Freshly isolated CD150^+^CD48^-^CD41^-^Flt3^-^CD34^-^ LSK cells (Fresh HSCs), or HSCs cultured 7 days in 4% nBSA (nBSA) or in 4% FA-free BSA supplemented with 200μg/mL each of palmitate and oleate (Reconstituted BSA) under 1% O2 conditions were subjected to cDNA microarray analysis. Cytokines (3ng/mL SCF and 0.1ng/mL TPO) were added to each culture condition. n = 1 for fresh HSCs, n = 1 for 4% nBSA, and n = 2 for reconstituted BSA samples. (**B**) Scatter plot of cDNA microarray data comparing samples shown in (**A**). Fatty acid or cholesterol synthesis genes are highlighted in red. Correlation coefficient *r* is noted. (**C**) Gene set enrichment analysis (GSEA) of cDNA microarray data shown in (**A**). FDR, false discovery rate. NES, normalized enrichment score. (**D**-**G**) Single-cell qPCR analysis (sc-qPCR). (**D**) Experimental design. Freshly isolated CD150^+^CD48^-^ CD41^-^Flt3^-^CD34^-^ LSK cells (Fresh HSCs), or HSCs cultured 7 days in 4% nBSA were subjected to sc-qPCR analysis. Cytokines (3ng/mL SCF and 0.1ng/mL TPO) were added to the culture media. After data filtering, 46 cells from the fresh HSC sample and 42 from the 4% nBSA sample remained. Data are from n = 1 experiment. (**E**) Principal component analysis of sc-qPCR data. The first two principal components are shown. (**F**) Violin plots showing expression levels of HSC-related genes. (**G**) Correlation coefficient matrix with hierarchical clustering of differentially-expressed genes between fresh and nBSA samples. sc-qPCR data from both groups were included in the analysis. (**H**) The effect of fatty acid or cholesterol supplementation on levels of differentially-expressed genes observed in either cDNA microarray or sc-qPCR was assessed by qPCR analysis of fresh and cultured HSCs. Freshly isolated CD150^+^CD48^-^Flt3^-^ LSK cells (Fresh HSCs), or HSCs cultured 7 days with 4% nBSA supplemented with the fatty acid mixture at indicated concentrations with or without 20μg/mL cholesterol and 20μg/mL palmitoyl phosphatidylcholine. The fatty acid mixture was as defined in Figure 2C and contained 3ng/mL SCF and 0.1ng/mL TPO. Shown are means±SD with n=4 technical replicates from 1 experiment. P values were calculated by two-way ANOVA to test effects of fatty acid (*P*_FA_) and cholesterol (*P*_chol_). The Fresh sample was omitted from ANOVA. (**I**) As in (**H**), effects of insulin concentration on cholesterol on expression of indicated genes were examined by qPCR. Freshly isolated CD150^+^CD48^-^Flt3^-^ LSK cells (Fresh HSCs), or HSCs cultured 7 days with 4% nBSA supplemented with the fatty acid mixture (as in (**I**)) at 1000μg/mL, with or without 20μg/mL cholesterol. Media contained 3ng/mL SCF and 0.1ng/mL TPO. Insulin was added at indicated concentrations. Shown are means±SD with n=4 technical replicates from 1 experiment. Indicated P values were calculated by two-way ANOVA. The fresh sample and samples in which no signal was detected were omitted from ANOVA.

We next assessed heterogeneity of gene expression in either fresh or cultured HSCs at the single cell level by performing single-cell qPCR (sc-qPCR) analysis (Figure 5D). Principal component analysis (PCA) showed that cultured HSCs skewed in both PC1-positive and PC2-positive directions in a pattern overlapping that of fresh HSCs (Figure 5E). Expression levels of HSC markers, including Ctnnal1 (Acar et al., 2015), Vwf (Sanjuan-Pla et al., 2013), and Gpr56 (Solaimani Kartalaei et al., 2015) or the representative HSC transcription factors Gata2 (de Pater et al., 2013), Pbx1 (Ficara et al., 2008), Tal1 (Shivdasani et al., 1995), and Erg (Loughran et al., 2008), were all high in cultured HSCs, while Fgd5 (Gazit et al., 2014), Procr (Kent et al., 2009), and Evi1 (Kataoka et al., 2011) expression showed cell-to-cell variability (Figures 5F and S5C), which caused a skew in the principal component space. Among differentiation markers, megakaryocytic genes including Mpl, Epor, CD9, F2r, and Itga2b (encoding CD41) were upregulated in the cultured HSCs, consistent with the microarray analysis (Figures S5B and S5D). On the other hand, lymphoid and myeloid differentiation-related genes, such as Flt3 and Csf3r (encoding G-CSF receptor), were downregulated in cultured HSCs (Figure S5C and S5D). Fatty-acid and cholesterol synthesis genes remained highly expressed in a fraction of cells, while genes encoding enzymes catalyzing fatty-acid oxidation were unchanged (Figures S5C and S5D). A correlation coefficient matrix with hierarchical clustering revealed that some HSC-related genes expressed in fresh HSCs, such as Egr1, Fgd5, Evi1, and Procr, formed a cluster with the myeloid-, and lymphoid-related genes Csf3r, Irf8, Flt3, and Ikzf1, whereas in cultured cells HSC-related Vwf, Ctnnal1, and Pbx1 formed a cluster with megakaryocytic genes as well as fatty acid synthesis genes (Figures 5G and S5E).

### Insulin induces a megakaryocytic differentiation program *in vitro*

We next modified the HSC culture medium as a means to establish gene expression patterns more closely resembling those seen in fresh HSCs. Further addition of fatty acids or fatty acids plus cholesterol to 4% nBSA medium dose-dependently downregulated Scd1 and Pmvk expression to levels seen in fresh HSCs, while the megakaryocyte genes or Evi1 or Flt3 remained unchanged (Figure 5H). The medium used in our experiments, SF-O3, contains supraphysiologically high insulin levels (∼10000-fold higher than serum concentration), which potentially activates AKT signaling and alters gene expression activating the IGF1 receptor. Decreasing insulin levels in maintenance medium downregulated expression of Gata1 and Sdpr, suggesting that insulin induces a megakaryocytic program (Figure 5I). In support of this idea, megakaryocyte number was positively correlated with insulin levels in the medium (Figure S5F). Interestingly, CD41 expression was inversely correlated with insulin concentration (Figure S5F). Addition of cholesterol to maintenance medium moderately increased the number of phenotypic HSCs at higher insulin concentrations, based on analysis of Sca-1 (Figure S5F). Both Evi1 and Flt3 expression was downregulated in any condition tested, although the mechanism remains unknown (Figures 5H-I, and S5F). Altogether, higher concentrations of fatty acid and cholesterol and reduced insulin levels altered gene expression patterns in cultured HSCs in a manner more closely resembling those seen in fresh HSCs, with the exception in Evi1 or Flt3 expression.

### Low cytokine levels and hypoxia increase HSC dependence on extrinsic fatty acids

As noted, cytokine concentration is a major quiescence/differentiation fate-determinant for HSCs *in vitro*. Since activity of cytokine target genes is strictly regulated in HSCs *in vivo* (Lee et al., 2010; Seita et al., 2007), we examined phosphorylation of SCF and TPO targets using intracellular flow cytometry. Targets analyzed were S6, ERK1/2, and STAT5, which are representative effectors of AKT, MAPK, and JAK/STAT signaling, respectively. To do so, we treated HSPC fractions from freshly isolated whole bone marrow cells with various combinations of SCF (0.33-81ng/mL, or none) and TPO (0.5-121.5ng/mL, or none) or with none for 10 to 30 minutes (Figures 6A and S6A-B). Relevant to signaling, HSCs are responsive to both SCF and TPO, whereas MPP cells respond primarily to SCF, reflecting HSC-specific expression of the TPO receptor c-Mpl (Buza-Vidas et al., 2006; Yoshihara et al., 2007). ERK phosphorylation increased with SCF concentration in each HSPC fraction, whereas S6 was more highly phosphorylated in HSCs than in MPPs in response to either SCF or TPO. In maintenance conditions (SCF 3ng/mL + TPO 0.1ng/mL; marked by an X in Figure 6A), both S6 and ERK were partially phosphorylated in all HSPC fractions. Moreover, in maintenance conditions, STAT5 phosphorylation in HSCs or MPPs was comparable and lower than that seen in HSCs treated with a high concentration (>13.5ng/mL) of TPO (Figures 6A and S6A-B). These results indicate that minimal activation of ERK and AKT signaling is required for HSC maintenance *in vitro*.

**Figure 6.**
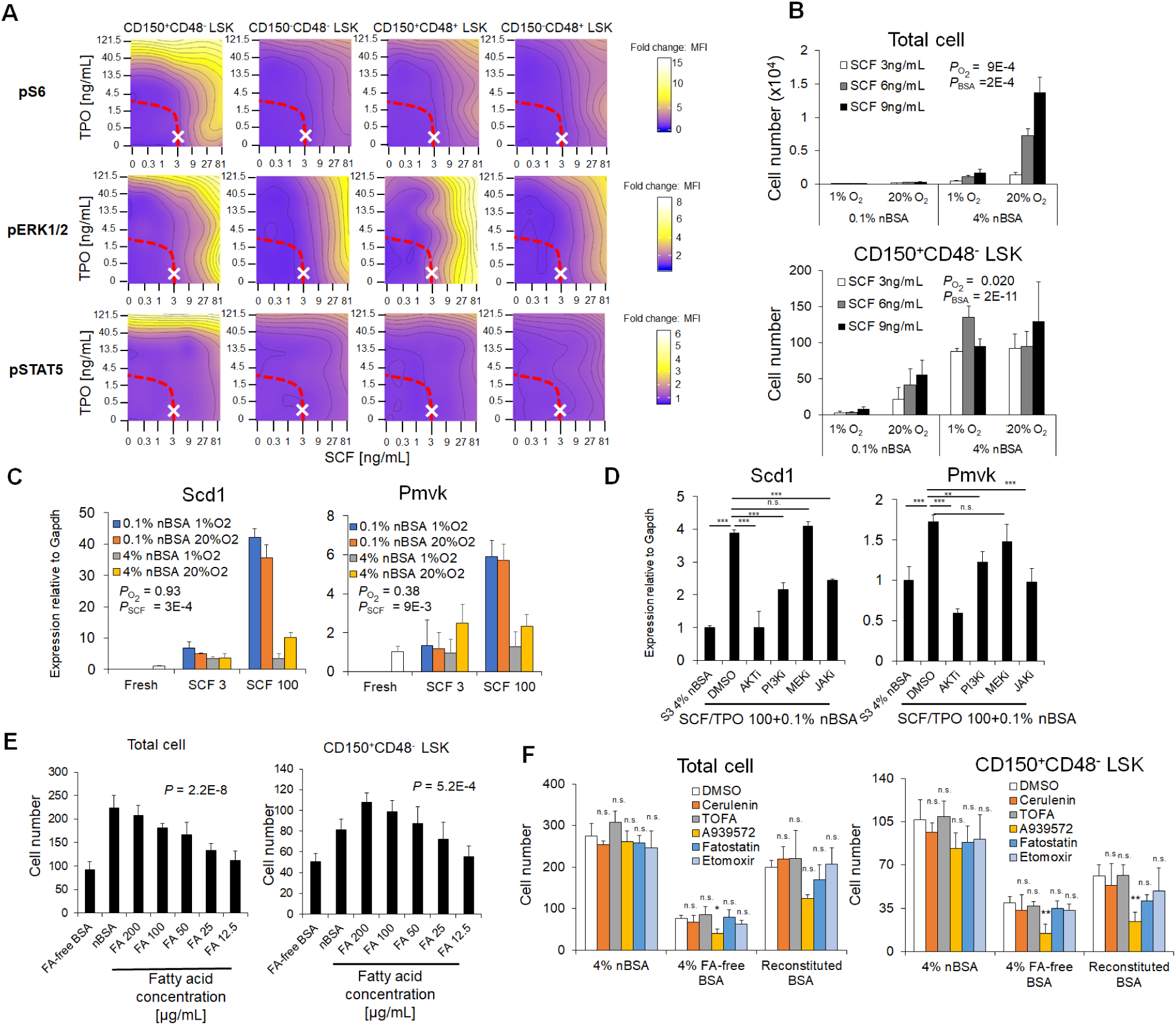
Low cytokine and hypoxic conditions enhance HSC dependence on exogenous fatty acids. (**A**) S6, ERK1/2, and STAT5 phosphorylation status in hematpoietic stem/progenitor fractions, as assessed by intracellular flow cytometry following exposure to SCF (0 to 81ng/mL) and/or TPO (0 to 121.5ng/mL). Dashed red line indicates the HSC maintenance border as shown in Figure 3D, and “X” marks 3ng/mL SCF and 0.1ng/mL TPO (n = 2 from 2 independent experiments for pS6, and 1 experiment for pERK1/2 and for pSTAT5). (**B**) HSC dependence on nBSA at low cytokine and hypoxic conditions. CD150^+^CD41^-^CD48^-^Flt3^-^CD34^-^ LSK cells (250) were cultured 7 days in indicated conditions and either total cell number (top) or the number of CD150^+^CD48^-^ LSK cells (bottom) was determined. All conditions contained 0.1ng/mL TPO (mean ± SD, n = 3). P values were calculated by two-way ANOVA to examine the influence of oxygen (*P*_O2_: 1% O_2_ or 20% O_2_) and nBSA (*P*_BSA_: 0.1% BSA or 4% BSA). (**C**) qPCR analysis of fatty acid (Scd1) and cholesterol (Pmvk) biosynthesis-related mRNAs after 16 hours of culture of CD150^+^CD41^-^CD48^-^Flt3^-^CD34^-^ LSK cells at 3ng/mL (S3) or 100ng/mL (S100) SCF in the presence or absence of 4% nBSA. Values were normalized to Gapdh expression and are shown as fold-induction relative to levels detected in freshly isolated (Fresh) CD150^+^CD41^-^CD48^-^Flt3^-^CD34^-^ LSK cells. Culture medium was DMEM/F12 lacking TPO, insulin, sodium selenite, and transferrin (mean ± SD, n = 4 technical replicate). P values were calculated by two-way ANOVA to examine the influence of oxygen (*P*_O2_: 1% O_2_ or 20% O_2_) and SCF (*P*_SCF_: S3 or S100). (**D**) qPCR analysis as in **(C)** but in the presence of kinase inhibitors. Values were normalized to Gapdh and are expressed as fold-induction compared to levels detected in 3ng/mL SCF plus 4% nBSA in 20% O_2_. Culture medium was DMEM/F12 lacking TPO, insulin, sodium selenite, and transferrin (mean ± SD, n = 4 technical replicates). AKTi, 1μM Triciribine. PI3Ki, 1μM PI-103. MEKi, 10μM MEK inhibitor I. JAKi, 1μM Jak inhibitor I. \**P* < 0.05, \*\**P* < 0.01, and \*\*\**P* < 0.001. n.s., not significant by Tukey-Kramer multiple comparison test. (**E**) Total cell number (left) and the number of CD150^+^CD48^-^ LSK cells (right) after 7 days culture of 300 CD150^+^CD41^-^CD48^-^Flt3^-^CD34^-^ LSK cells in either 4% FA-free BSA, 4% nBSA, or 4% reconstituted (200ug/mL each, sodium palmitate and sodium oleate) BSA under 1% O_2_ conditions. Shown are means ± SD, with n = 4. P values were calculated by one-way ANOVA to examine the influence of BSA or fatty acid concentration. (**F**) Total cell number (left) and the number of CD150^+^CD48^-^ LSK cells (right) after 8 days of culture of 300 CD150^+^CD41^-^CD48^-^Flt3^-^CD34^-^ LSK cells with either 4% nBSA, 4% FA-free BSA, or 4% reconstituted BSA under 1% O_2_ conditions in the presence of fatty acid synthesis inhibitors (5μM Cerulenin,1μM TOFA, 1μM A939572, or 20μM Fatostatin) or a fatty acid oxidation inhibitor (10μM Etomoxir) (mean ± SD, n = 4). \**P* < 0.05, \*\**P* < 0.01, and \*\*\**P* < 0.001. n.s., not significant compared with DMSO samples among all BSA samples, based on Tukey-Kramer multiple comparison test.

Next, we examined the nBSA requirement for HSC survival at lower cytokine concentrations (3-9ng/mL SCF) under hypoxia or normoxia. In 0.1% nBSA medium, HSCs did not survive at any SCF concentration, and the effect was exacerbated under 1% O_2_ conditions (Figure 6B). To test whether cytokine concentration or oxygen partial pressure positively regulates expression levels of genes functioning in fatty acid or cholesterol synthesis, we performed qPCR using fresh HSCs or HSCs cultured 16 hours in either 3ng/mL or 100ng/mL SCF, under 1% or 20% O2 conditions. Upregulation of Scd1 or Pmvk, which had been seen in HSCs cultured in 100ng/mL SCF, was suppressed at 3ng/mL SCF, independently of oxygen level. These observations may account for the requirement for higher nBSA levels under low cytokine conditions (Figure 6C). To identify signaling pathways relevant to fatty acid synthesis, HSCs cultured in 0.1% nBSA medium were treated with small molecule inhibitors of downstream cytokine targets for 16 hours and evaluated for Scd1 and Pmvk expression. Treatment with AKT, PI3K, or JAK inhibitors decreased expression levels of Scd1 and Pmvk, while a MEK inhibitor did not. Thus, PI3K/AKT and JAK signaling, but not that of MEK/ERK, likely mediates upregulated expression of fatty-acid and cholesterol synthesis genes (Figure 6D). We then evaluated HSC survival in media containing reconstituted BSA, the presence of reconstituted BSA dose-dependently rescued HSC survival (Figure 6E), a finding confirmed in transplantation analysis (Figures S4K-L). Inhibition of fatty acid synthesis or β-oxidation did not alter the number of HSCs cultured with 4% nBSA or 4% reconstituted BSA, indicating minimal dependence on these intrinsic pathways *in vitro* (Figure 6F). Of note, AKT inhibition in maintenance conditions in the presence of nBSA significantly decreased HSC number, suggesting that a dose determines whether AKT has a positive or negative effect on HSC maintenance (Figure S6C). Collectively, our findings suggest that HSCs incorporate extrinsic fatty acids for maintenance at lower cytokine levels and in hypoxia to compensate for compromised fatty acid biosynthesis.

### Human HSCs require both fatty acid and cholesterol for maintenance

We next assessed our maintenance conditions in cultures of human HSCs. First, we created a cytokine response map as we had in murine HSCs (Figure 3) using adult human bone marrow CD34^+^CD38^-^ CD90^+^CD45RA^-^ cells cultured with various SCF (0.3-81ng/mL, or none) and TPO (0.02-62.5ng/mL, or none) combinations under 1% O_2_ and 4% nBSA conditions. Like murine HSCs, human HSC proliferation paralleled cytokine concentration (Figure 7A). One striking difference, however, was that human HSCs depended more on TPO, with few cells remaining after a 7-day culture in TPO levels <1ng/mL (Figure 7A). Addition of FLT3L slightly rescued SCF and TPO dependence (data not shown). The number of total cells or CD34^+^CD38^-^ HSPCs was maintained at either SCF 3ng/mL + TPO 3ng/mL or SCF 1ng/mL +TPO 100ng/mL (Figure 7B). Human HSCs cultured under low cytokine levels and hypoxic conditions required nBSA for survival (Figure 7C).

**Figure 7.**
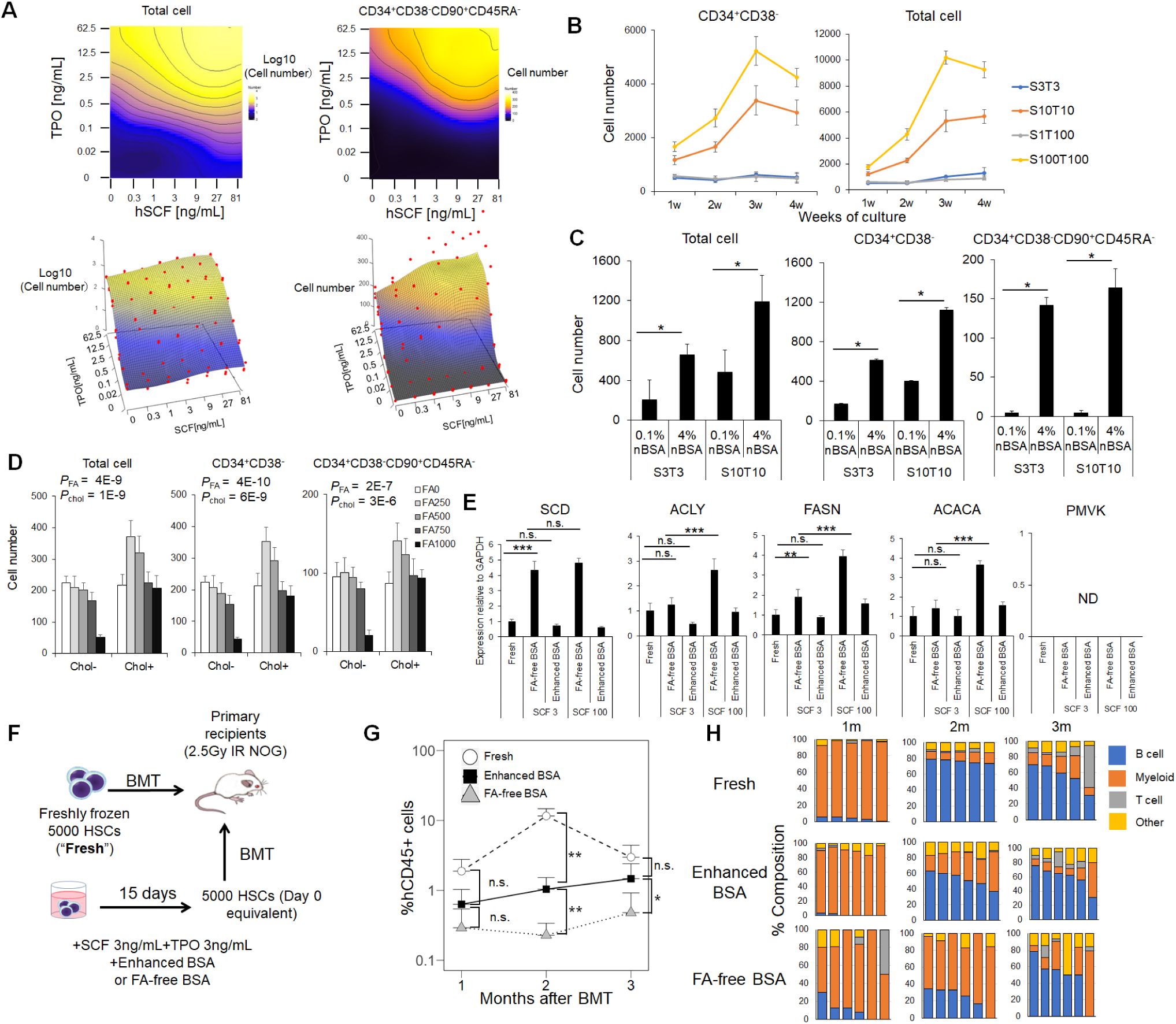
Human HSCs require both fatty acid and cholesterol for maintenance. (**A**) Contour plots of cell numbers after 7 days of culture (input: 600 CD34^+^CD38^-^CD90^+^CD45RA^-^ cells from a human bone marrow CD34^+^ sample) in SF-O3 + 4% nBSA and 1% O_2_ in 49 (7 x 7) different concentrations of SCF (0 to 81 ng/mL) and/or TPO (0 to 62.5ng/mL). Top panels show total cell number (left) and absolute number of phenotypic HSCs (CD34^+^CD38^-^CD90^+^CD45RA^-^ cells) (right). In bottom panels, surfaces predicted from data points observed in corresponding top panels are shown. (n = 3 wells per each condition from 1 experiment). (**B**) Total cell number (left) and the number of CD34^+^CD38^-^ cells after 1, 2, 3, or 4 weeks culture of 600 CD34^+^CD38^-^CD90^+^CD45RA^-^ cells with indicated cytokines. Medium was SF-O3 plus 4% nBSA, and cells were cultured in 1% O_2_ (mean ± SD, n = 4 for each group). (**C**) The effect of BSA and cytokine concentration on human HSCs. Cell number (total, CD34^+^CD38^-^, or CD34^+^CD38^-^CD90^+^CD45RA^-^ cells) was determined after 7 days of culture of 600 CD34^+^CD38^-^ CD90^+^CD45RA^-^ cells. Medium was SF-O3 plus 4% nBSA and SCF and TPO at 3ng/mL each, and cells were cultured in 1% O_2_ (mean ± SD, n = 4 for each group). \**P* < 0.05, \*\**P* < 0.01, and \*\*\**P* < 0.001. n.s., not significant by Tukey-Kramer multiple comparison test. (**D**) Effects of cholesterol and fatty acids on human HSCs. Cell number (total, CD34^+^CD38^-^, or CD34^+^CD38^-^ CD90^+^CD45RA^-^ cells) after 7 days of culture of 600 HSCs was determined. Medium was SF-O3 plus 4% nBSA with or without either the fatty acid mixture (palmitate, oleate, linoleate, and stearate at a 4:3:2:1 ratio) at indicated concentrations or 20μg/mL cholesterol. Cytokines were 3ng/mL SCF and 3ng/mL TPO. Cells were cultured in SF-O3 medium in 1% O_2_ conditions (mean ± SD, n = 4 for each group). P values were calculated by two-way ANOVA to examine the influence of fatty acid and cholesterol. (**E**) Gene expression in fresh or cultured human CD34^+^CD38^-^CD90^+^CD45RA^-^ cells was examined by qPCR. Cells were cultured 40 hours in either SCF and TPO at 3ng/mL each (SCF3) or SCF and TPO at 100ng/mL (S100) supplemented with either 4% FA-free BSA or 4% enhanced BSA (palmitate, oleate, linoleate, and stearate at a 4:3:2:1 ratio) and 20μg/mL cholesterol). Cells were cultured in αMEM (without exogenous insulin) in 1% O_2_ conditions. Shown are means ± SD, n = 4 for each group. \**P* < 0.05, \*\**P* < 0.01, and \*\*\**P* < 0.001. n.s., not significant by Tukey-Kramer multiple comparison test. ND, not detected. (**F**) Experimental design for bone marrow transplant of either fresh or cultured human CD34^+^CD38^-^ CD90^+^CD45RA^-^ cells into irradiated (2.5Gy) immunocompromised mice. The latter were cultured for 15 days in the presence of 4% FA-free BSA or 4% enhanced BSA (as defined in (**D**)) plus 20μg/mL cholesterol and SCF and TPO at 3ng/mL each. Cells were cultured in αMEM (lacking insulin) under 1% O_2_ conditions. 5000 cells (day 0-equivalent) were transplanted. (**G**) Peripheral blood chimerism of human CD45^+^ cells (mean + SD, n = 5 for Fresh; n= 6 for ehnhanced BSA; and n = 6 for FA-free BSA). \**P* < 0.05, and \*\**P* < 0.01. n.s., not significant, based on Wilcoxon’s rank sum test. (**H**) Composition of each lineage (Myeloid, B-cell, T-cell, and others) within hCD45^+^ cells at indicated time points after transplantation. Each bar represents data from individual recipients.

Next we examined fatty acid or cholesterol requirements using human HSCs. Addition of fatty acid alone to cultures containing nBSA dose-dependently decreased HSC number, while further addition cholesterol rescued HSC numbers (Figure 7D), suggesting a greater dependence on cholesterol by human HSCs. To determine fatty acid and cholesterol levels optimal for human HSC survival, we added varying (100-400μg/mL, or none) levels of a fatty acid cocktail, either with or without cholesterol, to nBSA (Figure S7B) and evaluated effects of those BSA preparations versus FA-free BSA (Figure S7D) in HSC cultures containing SCF and TPO at 3ng/mL each. The highest yield of HSCs was seen following culture in 200μg/mL cocktail plus either 20 or 40μg/mL cholesterol. Only a small increase in total cell number was seen in the nBSA condition (Figure S7B). In FA-free BSA cultures, total numbers of HSCs greatly decreased (Figure S7D), indicating that fatty acids and cholesterol are required for human HSC culture in low cytokine conditions. We then cultured HSCs in media containing SCF and TPO (both at 3ng/mL) and 4% nBSA plus 200μg/mL fatty acids and 20μg/mL cholesterol in 1% O_2._ (Hereafter, we designate nBSA plus 200μg/mL fatty acids/20μg/mL cholesterol as “enhanced BSA”.) We observed that ∼90% of cells exhibited a CD34^+^CD38^-^ marker phenotype, and 40% exhibited a CD90^+^CD45RA^-^ phenotype after 7 days of culture (Figure S7C), indicating that human HSCs remain quiescent with at least partially an undifferentiated phenotypes. To determine whether insulin is required for human HSC survival, we cultured HSCs under several insulin concentrations for 7 days. The number of human HSCs remaining in culture increased with insulin concentration, suggesting that insulin is required for HSC survival (Figure S7E). We then used qPCR analysis for human HSCs to determine if low expression of fatty acid and cholesterol synthesis genes seen at lower cytokine ranges underlies dependence on fatty acid and cholesterol. Like murine HSCs, human HSCs showed lower expression of some of fatty acid synthesis genes (ACLY, FASN, ACACA) at 3ng/mL compared to 100ng/mL SCF conditions. Interestingly, we did not detect PMVK mRNA in human HSCs in any conditions tested, including under fatty-acid-free conditions, supporting the high dependence of human HSCs on extrinsic cholesterol (Figure 7E). Fatty acid synthesis inhibition at high cytokine (50ng/mL SCF, 50ng/mL TPO, and 50ng/mL FLT3L) and normoxic conditions antagonized cell proliferation in both 0.1% nBSA or 4% FA-free BSA conditions, although inhibitor effects differed among groups (Figure S7F). In high or low cytokine conditions, cell number was maintained in the 4% nBSA condition, confirming dependence on extrinsic fatty acids (Figure S7F). Interestingly, cerulenin and fatostatin inhibited proliferation and survival of human HSCs cultured in 10% serum (Figure S7F), an outcome observed in analysis of murine HSCs (Figures 1G and 2E). Finally we evaluated the *in vivo* function of cultured human HSCs by transplanting them into immunocompromised NOD/Shi-scid, IL-2Rγ knockout (NOG) mice. HSCs were transplanted immediately after sorting or cultured with 3ng/mL SCF +3ng/mL TPO either with 4% enhanced BSA or 4% FA-free BSA under 1% O_2_ conditions. Three months after transplantation, the enhanced BSA group contributed equally to peripheral blood chimerism, relative to fresh HSCs, and showed a balanced output of B cell, T cell, and myeloid lineages. By contrast, the FA-free group contributed less to the peripheral blood chimerism than did either fresh or enhanced BSA groups (Figures 7F and 7G), indicating that supplementation with fatty acid and cholesterol favors HSC maintenance in vitro (Figures 7F-H).

## DISCUSSION

Manipulation of HSCs *in vitro* requires technology to maintain them in an environment resembling bone marrow. Here, we devised a near-physiological condition to maintain quiescent HSCs *in vitro* with defined factors. Key features of our culture conditions were adequate supplementation of fatty acid (and cholesterol, in the case of human HSCs), a low cytokine concentration, and hypoxia. Proper metabolic activity plays crucial role in regulating HSC survival and function *in vivo*. We show here that metabolic changes induced by conventional HSC culture were minimized by exogenous fatty acid. Fatty acids serve to maintain cellular membrane properties (Currie et al., 2013) or signaling and in energy production through β-oxidation (Ito et al., 2012). Previously, Ito et al (2012) reported that treatment of HSCs with a β-oxidation inhibitor induced a loss of HSC maintenance. By contrast, here, we observed minimal effects of treatment of HSCs with β-oxidation inhibitor, suggesting that fatty acids provide essential membrane components or signaling factors but do not function in energy production in this context (Figure 6F). Ito et al. inhibited β-oxidation to promote *in vitro* HSC division under high cytokine conditions (50 ng/mL SCF, 50 ng/mL TPO) in which HSCs may require greater energy production. Thus, discrepancies between that study and ours may attributable to differences in culture conditions. We did not test *in vivo* inhibition of β-oxidation; however, we do not exclude the possibility that quiescent HSCs are more dependent on fatty acid oxidation than on fatty acid synthesis *in vivo*, given that expression of fatty acid synthesis genes are suppressed in freshly-isolated HSCs (Figure 1A).

Recently, liver-derived TPO was shown to extrinsically regulate HSCs outside bone marrow (Decker et al., 2018). Liver-derived circulating albumin may be another liver-derived molecule regulating HSC dynamics *in vivo*. Particularly in human HSCs, cholesterol was required for HSC maintenance, possibly due to lack of PMVK expression in human HSCs. Cholesterol regulates membrane homeostasis and protein modification and provides steroid analogues including bile acid, all required for HSC maintenance (Sigurdsson et al., 2016). Notably, cholesterol is required for HSCs, especially at high fatty acid levels (Figure 7D); thus HSC maintenance may require a proper fatty acid/cholesterol balance more than an absolute level cholesterol. Defined media used previously for HSC culture may not have contained sufficient levels of cholesterol to allow proper HSC maintenance.

Lower cytokine concentrations are critical to maintain HSCs in an immature state but may insufficient for de novo biosynthesis of essential factors such as fatty acids (as shown here), possibly due to suppressed expression of biosynthetic enzymes. These results are consistent with accumulating finding showing that AKT activation perturbs HSC maintenance and that quiescent HSCs are metabolically inactive (Chandel et al., 2016). Average oxygen concentration is as low as 1.3% around the sinusoid of bone marrow (Spencer et al., 2014). That hypoxic environment plays an important role in HSC quiescence, glycolytic metabolism, and retention in the niche (Cipolleschi et al., 1993; Takubo et al., 2010). Hypoxia reportedly maintains proper function and decreases cycling of cultured human HSCs (Danet et al., 2003). By combining lower cytokine and higher fatty acid concentrations with hypoxia, we enhanced this activity and established a low proliferation state *in vitro*. In our system, hypoxia suppressed HSC differentiation by modulating responses to cytokines: HSC maintenance was enhanced by SCF, while megakaryocyte differentiation was suppressed by TPO. In addition, hypoxia conferred nBSA dependence on HSCs, especially in low SCF conditions. In addition to physiologically low oxygen tension in the bone marrow space (Spencer et al., 2014), free SCF concentrations in bone marrow fluid are also kept low (Asik et al., 2003). Also, albumin concentrations in serum are approximately 3.5-5.0% w/v. Thus, HSC nutrient and environmental requirements as defined here closely follow physiological conditions of oxygen, cytokines, and fatty acid-bound albumin, and our system allows assessment of factors that regulate HSC cell cycle and/or differentiation status.

Although we achieved 1-month maintenance of functional HSCs *in vitro*, our system does not fully recapitulate HSC conditions *in vivo*. For example, we observed 1) gradual loss of HSC markers that normally show robust expression *in vivo*, among them, CD150 and Evi1, 2) lower chimerism, especially after primary transplantation with lower lymphoid potential, and 3) higher expression of genes marking aging HSCs, such as CD41. These differences are likely due to conditions or factors missing in the current culture condition. Cytokine levels in the bone marrow microenvironment depend on many variables, including local concentration, distances from cytokine sources (which change over time due to HSC movement), diffusion coefficients of each molecule, and cytokine sequestration by soluble or membrane-bound receptors. All of these factors make defining cytokine conditions in the HSC environment *in vivo* challenging, and fine temporal manipulation of extracellular cytokine concentration may improve HSC maintenance in culture. Nutrients other than fatty acids, such as glucose (Takubo et al., 2013), amino acids (Taya et al., 2016), and BSA contaminants (Ieyasu et al., 2017) may also modulate HSC fate, as may lipid components. Comprehensive nutrient and cytokine optimization as well as more accurate measurement of extrinsic factors in the native bone marrow will facilitate reconstitution of the bone marrow microenvironment *ex vivo*.

The system defined here has numerous potential benefits for HSC research and engineering. A more efficient culture system will allow (1) better analysis of HSC behavior in *ex vivo* conditions, (2) detailed characterization of HSC-specific effects of cytokines or chemical compounds, and (3) improved engineering of HSCs for gene therapy by minimizing differentiation or loss of stem cell capacity. These merits are equally beneficial to the capture of HSCs generated from pluripotent stem cells. The new system also provides insight into HSPC behavior *in vivo*. HSCs are by nature slow-cycling in the absence of factors except for SCF and TPO, and some conditions preferentially activate the MPP4 cell cycle, raising the possibility that a specialized niche favoring HSC rather than MPP quiescence is not necessary. HSCs and MPPs may behave differently under identical conditions in terms of cell cycle activation and self-renewal capacity. Also, this culture system enables analysis of HSC heterogeneity or of gradual changes in HSC composition that occur within the HSC pool. Overall, the *ex vivo* system described here will further facilitates analyses of HSC function and dynamics and enable more efficient manipulation of HSCs.

## Supporting information

## ACKNOWLEDGEMENTS

We thank all members of the Takubo laboratory for indispensable support; M. Haraguchi and S. Tamaki for technical support and laboratory management; and E. Lamar for preparation of the manuscript. KT was supported in part by KAKENHI Grants from MEXT/JSPS (26115005, 18H02845, 18K19570, 26115001, 15K21751), a grant of the National Center for Global Health and Medicine (26-001, 29-2007), AMED-CREST (JP18gm0710010), an AMED grant for Realization of Regenerative Medicine (JP18bm0704011) and grants from the Japan Leukemia Research Fund, the Japan Rheumatism Foundation, the Takeda Science Foundation, the Senshin Medical Research Foundation, and the Japanese Society for Hematology. HK was supported in part by a KAKENHI Grant (17K16200), a grant of the National Center for Global Health and Medicine (29-1015) and a grant from the Uehara Memorial Foundation. HS was supported in part by AMED-CREST (JP18gm0910011 and JP18gm0710002). MS was the leader of JST ERATO Suematsu Gas Biology until March 2015, which provided infrastructure for multi-photon laser confocal microscopy, essential to accomplish the aim of this current study.

## AUTHOR CONTRIBUTIONS

H.K., T.M., A.O., F.H., S.W., and T.H-Y. performed the study and analyzed data; G.N., D.H., H.S., F.A., Y.K., M.S., and T.S. provided scientific advice and materials. H.K. and K.T. wrote the manuscript. K.T. conceived the project and supervised the research.

## Methods

### CONTACT FOR REAGENT AND RESOURCE SHARING

Further information and requests for resources and reagents should be directed to and will be fulfilled by the Lead Contact, Keiyo Takubo

### EXPERIMENTAL MODELS AND SUBJECT DETAILS

#### Mice

C57BL/6J mice (8-14 week-old, purchased from SLC Japan or CLEA Japan) were used in all experiments, unless otherwise stated. C57BL/6-Ly5.1 congenic mice purchased from CLEA Japan were used for competitive repopulation assays. Ubc-GFP reporter mice (Schaefer et al., 2001) were purchased from The Jackson Laboratory and used as donors in reconstitution assays to examine platelet and red blood cell frequency. NOD/Shi-scid,IL-2RγKO Jic (NOG) mice were purchased from In-Vivo Science Inc. and used for human cell reconstitution assays. All mice were bred in the animal facility at the National Center for Global Health and Medicine under specific pathogen-free (SPF) conditions and fed ad-libitum. Mice were euthanized by cervical dislocation. Animal experiments were approved by the National Center for Global Health and Medicine. Both male and female mice were used in experiments.

#### Human CD34^+^ bone marrow cells

Human CD34^+^ bone marrow cells were purchased from Lonza and stored in liquid nitrogen until use. Frozen cells were thawed in vials in a 37°C water bath and transferred to a 15mL tube. Medium (SF-O3 or αMEM or DMEM/F12) containing 10 % FCS plus 2U/mL DNaseI (Sigma Aldrich) was slowly added to the suspension while swirling gently to fill the tube. The suspension was centrifuged at 200 x g for 15 minutes at room temperature. Supernatants were aspirated and the wash step repeated. An antibody cocktail (50μL of PBS + 2%FCS plus 10μL of anti-CD34-FITC, 2μL of anti-CD38-PerCP-Cy5.5, 5μL of anti-CD90-PE-Cy7, and 10μL of anti-CD45RA-PE) was added to the suspension and kept on ice for 30 minutes. Cells were washed once with PBS+2% FCS and resuspended in PBS+2%FCS with 0.1% PI. Cells were sorted into SF-O3 or αMEM containing 4% BSA + 55μM 2-ME by FACSAria II.

### METHOD DETAILS

#### Cell preparation

Mouse bone marrow cells were isolated from two femurs and tibiae. Femurs and tibiae were flushed with PBS+2% FCS using a 21-gauge needle (Terumo) and a 10mL syringe (Terumo) to collect the bone marrow plug. The plug was dispersed by refluxing through the needle and the suspension centrifuged 680 x *g* for 5 minutes at 4°C. Cells were then lysed with lysis buffer (0.17M NH_4_Cl, 1mM EDTA, 10mM NaHCO_3_) at room temperature for 5 minutes, washed with 2 volumes PBS+2%FCS, and centrifuged at 680 x *g* for 5 minutes at 4°C. Cells were resuspended in PBS + 2%FCS and filtered through 40μm nylon mesh (BD Biosciences). Cells were again centrifuged 680 x *g* for 5 minutes at 4°C and treated with anti-CD16/32 antibody for Fc-receptor block (2μL/mouse) for 5 minutes at 4°C followed by addition of anti-c-Kit magnetic beads (Miltenyi) at a 1/5 volume/volume ratio for 15 minutes at 4°C. After removing the antibody with two PBS+2%FCS washes, c-Kit-positive cells were isolated using Auto-MACS Pro (Miltenyi) using the Possel-S program. Isolated cells were once centrifuged 340 x *g* for 5 minutes and stained with antibodies for flow cytometry.

#### Cell culture

SF-O3 medium (Sanko Junyaku) is a mixture of RPMI1640, Dulbecco’s MEM, and Ham’s F-12 medium at a 2:1:1 ratio with added cytidine, glutathione, para-amino benzoic acid, and cholesterol (a formula suggested by Shin-ichi Nishikawa, personal communication) supplemented with recombinant-human insulin, recombinant-human transferrin, sodium selenite, ethanolamine, HEPES, and sodium bicarbonate 2.2g/L. We further added 0.1% v/v of BSA by diluting a 10% BSA concentrate (Sanko Junyaku) or adding 0.1% w/v BSA powder (Sigma Aldrich). Additional BSA powder or fatty-acid-free BSA (Sigma Aldrich) was directly dissolved in the medium at indicated concentrations and filtered using 0.22μm filter. 2-ME was added before filtration at final concentration of 55μM. SCF, TPO, and other cytokines or reagents were added to the filtered media.

For other culture media (αMEM or DMEM/F12 medium), we added human recombinant insulin (Nacalai Tesque) before filtration at indicated final concentrations. Sorted murine or human cells were centrifuged at 340 x *g* for 5 min and supernatants discarded. Media was added to dilute cells to the desired number of cells per well in 10μL, which was dispensed into 200μL of culture media. Culture conditions were either 1% O_2_ + 5% CO_2_ or 20% O_2_ + 5% CO_2_ at 37°C in appropriate humidified incubators. For seven-day cultures, medium remained unchanged. For longer periods (11-31 days), half of the medium was changed twice a week.

#### Bone marrow transplant

For donor cells, HSCs (CD150^+^ CD41^-^ CD48^-^ CD34^-^ Flt3^-^ LSKs) from C57BL/6-Ly5.2 mice or HSCs (CD150^+^CD41^-^CD48^-^Flt3^-^ LSKs) from Ubc-GFP mice or cultured HSCs, together with 4×10^5^ BMMNCs from untreated C57BL/6-Ly5.1 or Ly5.2 mice, were used. For recipients, C57BL/6-Ly5.1 or Ly5.2 congenic mice were used. Donor cells were transplanted retro-orbitally into recipients that had been lethally-irradiated (9.5Gy using MBR-1520R (Hitachi Power Solutions), 125kV 10mA, 0.5mm Al, 0.2mm Cu filter). At 1, 2, 3, and 4 months after BMT, peripheral blood was collected and the percentage of donor-derived cells and their differentiation status determined by MACSQuant. Forty to eighty μL of peripheral blood was sampled from the retro-orbital plexus using heparinized glass capillary tubes (Drummond Scientific) and was suspended in 1mL of PBS+heparin. For platelet and red blood cell (RBC) analysis, 5μL of blood suspension was mixed with 50μL staining solution (PBS + 2% FCS with Ter-119-PerCP-Cy5.5 (1:100), and anti-CD41-PE(1:100)) for 30 minutes at 4°C, washed once and analyzed by MACSQuant. For white blood cell analysis, the blood suspension was centrifuged at 340 x *g* for 3 minutes. The supernatant was discarded and the pellet was resuspended into 1mL of PBS + 1.2% w/v dextran (200kDa, Nacalai Tesque) for 45 minutes at room temperature. The supernatant then was centrifuged at 340 x *g* for 3 minutes, and pellets were resuspended in 0.17M NH_4_Cl solution to lyse residual RBCs for 5 to 10 minutes until the suspension became clear. Cells were resuspended in 50μL PBS +2%FCS with 0.3μL Fc-block. Surface antigen staining was performed using the following antibody panel: Gr1-PE-Cy7, Mac-1-PE-Cy7, B220-APC, CD4-PerCP-Cy5.5, CD8a-PerCP-Cy5.5, CD45.1-PE, CD45.2-FITC. All antibodies were 0.3μL per sample. The frequency (%) of donor-derived cells was calculated as follows:

100 x Donor-derived (GFP^+^ or Ly5.2^+^Ly5.1^-^) cells (%) / (Donor-derived cells (%) + Competitor- or recipient-derived (Ly5.2^-^Ly5.1^+^ or GFP^-^) cells (%))

Myeloid cells, B cells, T cells, RBCs, and platelets were marked by Gr-1^+^ or Mac-1^+^, B220^+^, CD4^+^ or CD8^+^, CD41^-^ Ter119^+^, and CD41^+^ Ter119^-^, respectively. Total cell chimerism represents the frequency of donor-derived Ly5.2^+^Ly5.1^-^ or GFP^+^ cells over the frequency of Ly5.2^-^Ly5.1^+^ or GFP^-^ cells in mononuclear or unlysed cells. At 4 months after BMT, the frequency of donor-derived cells in bone marrow was determined using one femur and tibia per recipient. After counting bone marrow cells using an automated counter TC10 (Bio-Rad), equal volumes of cell suspensions (20-30% of total volume) from each recipient were pooled and 2 x 10^6^ of cells were resuspended in SF-O3 medium (0.1% BSA). The cell suspension (200μL/recipient) was transplanted by retro-orbital injection into lethally (9.5 Gy)-irradiated Ly5.1^+^ recipients with a 1mL syringe and 27-gauge needle. Remaining cells were stained to assess bone marrow chimerism. Anti-CD150-BV421, anti-CD48-PE, anti-Flt3-APC, anti-lineage (CD4, CD8a, Gr-1, Mac-1, Ter-119, B220)-PerCP-Cy5.5, anti-c-Kit-APC-Cy7, anti-Sca-1-PE-Cy7, anti-Ly5.1-Alexa-Fluor700, and anti-Ly5.2-FITC were used for surface antigen detection. All antibodies were 1μL per sample.

#### Bone marrow transplant of human HSCs into NOG mice

A total of 5000 HSCs (CD34^+^ CD38^-^ CD90^+^ CD45RA^-^) from frozen samples or the progeny of 5000 cultured HSCs (day 0 equivalent) were transplanted retro-orbitally into irradiated (2.5Gy) NOG mice. At 1, 2, and 3 months after BMT, peripheral blood was collected and the percentage of human cells and their differentiation status were determined by MACSQuant. After cell preparation as described above, cells were resuspended in 50μL PBS +2%FCS with 0.3μL anti-mouse Fc-block. Surface antigen staining was performed using the following antibody panel: mouse CD45-PE-Cy7, mouse Ter-119-PE-Cy7, human CD45-BV421, human CD13-PE, human CD33-PE, human CD19-APC, human CD3-APC-Cy7.

Myeloid cells, B cells, or T cells were determined as human CD45^+^ CD13/CD33^+^, human CD45^+^ CD19^+^, or human CD45^+^ CD3^+^ respectively. Total cell chimerism represents the frequency of PI human CD45^+^ murine CD45^-^ Ter-119^-^ cells over total PI^-^ mononuclear cells.

#### Fatty acid and cholesterol reconstitution

Fatty acid sodium salt (a combination of palmitate, oleate, linoleate, and stearate) was dissolved in methanol to a concentration of 4-20mg/mL and cholesterol (Tokyo Chemical Industry Co., Ltd.) was separately dissolved in methanol to a concentration of 4mg/mL in glass tubes (Maruemu Corporation). Solutions were mixed in glass tubes, air-dried, and then heated on water-bath at 50°C until methanol had evaporated. Medium containing 4% w/v BSA was directly added to the tube and sonicated until lipids were dissolved. Medium was then filtered using a 0.22μm filter (Millipore). 2-ME or insulin (if needed) was added just before filtration. Other reagents were added after filtration.

#### BSA

We used four types of BSA in culture media: native BSA (nBSA, Sigma Aldrich), fatty-acid-free BSA (FA-free BSA, Sigma Aldrich), FA-free BSA reconstituted with fatty acids (reconstituted BSA), and nBSA reconstituted with fatty acids and cholesterol (enhanced BSA). Fatty acid stock solutions used for reconstituted BSA were either 1:1 sodium palmitate and sodium oleate or 4:3:2:1 sodium palmitate, sodium oleate, sodium linoleate, and sodium stearate. Fatty acids used for enhanced BSA were 4:3:2:1 sodium palmitate, oleate, linoleate, and stearate. In that case, the final concentration of fatty acids was 400μg/mL and of cholesterol was 20μg/mL.

#### Apoptosis analysis

Cultured HSCs were stained using an Annexin V-PE Apoptosis Detection Kit (BD Biosciences), according to the manufacturer’s instructions. Cells were resuspended in 250μL of PBS+2%FCS +0.1%PI, and apoptotic cells (Annexin V^+^ PI^-^ cells) were detected by MACSQuant.

#### EdU incorporation assay

Cells were cultured 16 hours under indicated conditions, and 10mM EdU was then added to culture medium to a final concentration of 10μM and incubated at 37°C for 2 hours. Cells were permeabilized to detect intracellular EdU using a Click-iT EdU Alexa Fluor 647 Flow Cytometry Assay Kit (Thermo Fisher Scientific) and analyzed by MACSQuant (Miltenyi Biotec).

#### Intracellular flow cytometry for phosphorylated ERK1/2, S6, and STAT5

Murine BMMNCs were pooled from 4 mice per experiment. Cells were treated with 8μL (1:50) of Fc block for 10 minutes at 4°C. Since antigenicity of CD150 and Sca-1 is vulnerable to the fixation/permeabilization process, these markers were stained in advance. Cells were stained with 8μL (1:50) of CD150-BV421 antibody and 8μL (1:50) of Sca-1-Alexa700 antibody at 4°C for 30 minutes and then washed once in PBS + 2% FCS and resuspended in DMEM/F12 medium at 4 x10^7^cells/mL. 50μL of cell suspension was dispensed to each well of 96-well round bottom plates followed by addition of 50μL DMEM/F12 containing 2x concentrations of cytokines. Plates were incubated at 37°C, 5%CO_2_, and 20% O_2_ conditions for indicated times. Cells were then fixed for 10 minutes at 37°C by adding 100μL of Fixation buffer I (BD Biosciences, PBS with 4% paraformaldehyde) and then chilled immediately by transferring to ice-cold 1mL of PBS/2% FCS. Cells were centrifuged (600 x *g* for 3 minutes) and then permeabilized by addition of 500μL of ice-cold 90% methanol followed by incubation on ice for 30 minutes. After washing twice with 1mL PBS + 2% FCS (staining solution), cells were resuspended in 50µL staining solution containing anti-lineage marker (CD4, CD8a, B220, Ter-119, Gr-1, Mac-1)-PerCP-Cy5.5 (1:100), anti-c-Kit-APC-Cy7 (1:100), anti-CD48-PE (1:100), and either anti-phospho-ERK1/2-AlexaFluor488 antibody (1:5), or anti-phospho-AKT-AlexaFluor488 antibodies (1:5), or anti-phospho-STAT5-AlexaFluor488 antibody (1:5) for 1 hour at room temperature. Stained cells were washed once with 1mL staining medium and then resuspended in 250μL staining medium, and the samples were acquired by FACSAria II. Data were analyzed using FlowJo software.

#### Single-cell qPCR analysis

Fresh CD150^+^CD48^-^CD41^-^Flt3^-^CD34^-^ LSK cells (HSCs) from 10 mice were sorted into 500µL SF-O3 containing 4% w/v BSA +2-ME. 33000 HSCs were split into 2 tubes, one (fresh HSCs) subjected to reverse transcription and preamplification using Fluidigm C1 system (Fluidigm) and the other cultured 7 days in 2 wells of a 96-well round-bottom plate (cultured HSCs) followed by reverse transcription and preamplification. SF-O3 containing 4% w/v BSA + 55μM 2-ME with SCF 3ng/mL plus TPO 0.1ng/mL was used for culture medium. Reverse transcription and pre-amplification was performed according to the manufacturer’s instruction using an Ambion Single Cell-to-CT kit (Thermo Fisher Scientifc) and TaqMan PCR probes (Thermo Fisher Scientific). Cells were diluted in Suspension Reagent (Fluidigm) to ∼1000 cells/µL and loaded to C1-Single-Cell Auto Prep IFC for Preamp (5-10μm). After loading cells, IFC was observed microscopically to ensure that single cells were captured in each well using IN Cell Analyzer 6000 (GE Healthcare). Wells with 0 or >1 or shrunken cells were excluded. A total of 80 wells of 96 for fresh HSCs, and 62 of 96 wells for cultured HSCs were regarded as valid. Thermal cycling conditions in reverse transcription, and preamplification were as follows: 25°C for 10min, 42°C for 1hr, and 85°C for 5 min for reverse transcription, and 95°C for 10min, 18 cycles of 95°C for 15s and 60°C for 4min, and kept 4°C until harvest. Amplified cDNA (about 3.5μL) from each sample was diluted with 25μL C1 DNA Dilution Reagent (Fluidigm) and frozen at -30°C until use. 48 samples of cultured or Fresh HSCs were subjected to qPCR. For single-cell qPCR analysis, a Biomark system (Fluidigm) in combination with Fluidigm 96.96 Dynamic Array IFC was used. For assay mix, 2.5µL of TaqMan Gene Expression Assay (Thermo Fisher Scientific) and 2.5µL of Assay Loading Reagent (Fluidigm) were mixed for each gene. For the sample mix, 2.5µL of TaqMan Fast Universal PCR Master Mix (Thermo Fisher Scientific), 0.25µL of GE Sample Loading Reagent (Fluidigm), and 2.25µL of cDNA were mixed. Assay and sample mixes were loaded onto 96.96 Dynamic Array IFC, and PCR analysis was performed on BioMarkHD.

#### Flow cytometry and cell sorting

After selection of c-Kit+ cells using magnetic beads, the murine hematopoietic stem and progenitor fraction was labeled as follows. For staining of C57BL/6J mice, lineage marker (CD4, CD8a, Gr-1, Mac-1, Ter-119, B220)-PerCP-Cy5.5, c-Kit-APC-Cy7, Sca-1-PE-Cy7, CD150-PE, CD41-FITC, CD48-FITC, Flt3-APC, CD34-BV421 were used. For staining of Ubc-GFP mice, lineage marker (CD4, CD8a, Gr-1, Mac-1, Ter-119, B220)-PerCP-Cy5.5, c-Kit-APC-Cy7, Sca-1-PE-Cy7, CD150-BV421, CD41-PE, CD48-PE, Flt3-APC were used. All antibodies used were 0.5µL per mouse. Cells were resuspended in 0.5 to 2mL of PBS +2%FCS + 0.1%PI and sorted by FACSAria II into SF-O3 containing 4% w/v BSA or other medium (αMEM or DMEM/F12) with 4% w/v BSA. For murine cell experiments, when cultures were conducted in fatty-acid-free conditions, fatty-acid-free BSA was used for sorting medium. For human cell experiments, fatty-acid free BSA was not used for sorting medium due to low cell survival rate. Murine HSCs were defined as CD150^+^CD41^-^CD48^-^CD34^-^Flt3^-^LSK or CD150^+^CD41^-^CD48^-^LSK (Kiel et al., 2005) or CD150^+^CD48^-^LSK cells. Multipotent progenitors (MPPs) were subfractionated as MPP1s (CD150^-^CD41^-^CD48^-^Flt3^-^LSK), MPP2s (CD150^+^CD41/CD48^+^Flt3^-^LSK), MPP3s (CD150^-^CD41/CD48^+^Flt3^-^ LSK), or MPP4s (Flt3^+^LSK). Human HSCs were defined as CD34^+^CD38^-^CD90^+^CD45RA^-^. Data were analyzed using FlowJo^™^ software (Tree Star Inc.).

#### Intravital Imaging of BSA

Male mice (24–31g, 11–13 weeks old, non-fasting) were anesthetized via intraperitoneal injection of urethane (800mg/kg) and α-chloralose (80mg/kg), tracheotomized, and intubated with a handmade Y-shaped tube for mechanical ventilation. Animals were mechanically ventilated with a small-animal ventilator (MiniVent type 845, Harvard Apparatus) with 21% O_2_ at a tidal volume of 8μL/g and a respiratory rate of 120 breaths/min. The left femoral artery and vein were cannulated to monitor mean arterial pressure (MAP) and intravenous chemical administration, respectively. An arterial catheter connected to a pressure transducer (MP 150, BioPac Systems) was placed in the left femoral artery to continuously monitor MAP and heart rate. Rectal temperature was maintained at 37.0 ± 0.5 °C throughout the experiment using a heating pad (ATC-2000, World Precision Instruments). The head of each mouse was fixed in a stereotactic frame (SG-4 N, Narishige Scientific Instrument Lab) in the sphinx position. The skull bone was exposed by a midline skin incision. Images were acquired using a two-photon laser microscope (FV1000MPE, Olympus) attached to a mode-locked titanium-sapphire laser system (Chameleon Vision II, Coherent) that could achieve a 140-fs pulse width and an 80-MHz repetition rate. To visualize the bone marrow microvasculature, 500 kDa TRITC-dextran (0.2g/kg body weight, Merck) was injected. FITC-albumin (40mg/kg body weight, Merck) was administered intravenously to assess vascular leakage of albumin. FITC-albumin intensity was measured using Fluoview software (version FV10-ASW, Olympus).

#### Measurement of albumin in murine BM and serum

Forty to eighty μL of peripheral blood was sampled from the retro-orbital plexus <u>of mice</u> using heparinized glass capillary tubes (Drummond Scientific) and collected into BD Microtainer blood collection tubes (BD Biosciences). For BM extracellular fluid collection, one femur and one tibia were placed on the filter membrane of 10μL Barrier Pipette Tips (Sorrenson BioScience). The tip was cut to a 5mm and then placed in a 1.5mL tube and centrifuged 15000*g* for 2 minutes. Each 0.5μL of serum or BM extracellular fluid was diluted at 1:40000, 1:160000, and 1:640000 in PBS and analyzed using a mouse Albumin ELISA Kit (Abcam), according to the manufacturer’s instruction. Data from the 1:160000 dilution sample was considered valid as values were within the range of the calibration curve.

The serum was subjected to an ELISA using a Quantikine ELISA Kit (R&D Systems)

#### Measurement of fatty acid content of BSA

Fatty acid measurement was performed as described (Shimura et al., 2016). All fatty acids in 10mg samples of lyophilized native BSA (nBSA) or fatty-acid-free (FA-free) BSA were first derivatized to fatty acid methyl esters (FAMEs) using a FAME derivatization and purification kit (Nacalai Tesque), according to the manufacturer’s instructions. C23:0 (Supelco n-Tricosanoic acid, Sigma–Aldrich) was added to BSA samples prior to methylation as an internal standard. FAMEs were analyzed by a gas chromatograph with electron impact mass spectrometry (GCMS-QP2010 Ultra (Shimadzu), using a FAMEWAX capillary column (30 m × 0.25 mm I.D. × 0.25 μm) (Restek Corporation). The injection port temperature was set at 250°C. A 2-μl aliquot was injected in splitless mode. Column temperature was programmed as follows: initial temperature was held at 40°C for 2 min; increased at 20 to 140°C/min, 11 to 200°C/min, and 3°C to 240°C/min. It was then maintained at 240°C for 10 min. Helium served as carrier gas with a linear velocity of 45 cm/s. FAMEs were detected in the selected ion monitoring mode of the characteristic fragment ions (m/z 55, 67, 74, and 79) and quantified using peak areas of known amounts of FAMEs (Supelco 37 Component FAME Mix, Sigma–Aldrich) and C23:0 FAME.

#### MACSQuant analysis of cell number

Most (170µL) of the medium in wells of a 96-well plate was aspirated and samples were stained with 10μL of antibody cocktail for 30 minutes at 4°C. For murine experiments, antibodies used were anti-lineage markers (CD4, CD8a, Gr-1, Mac-1, B220, Ter-119)-PerCP-Cy5.5, anti-c-Kit-APC-Cy7, anti-Sca-1-PE-Cy7, anti-CD150-PE, anti-CD48-FITC, anti-CD41-APC. All antibodies used were 0.1μL/well. For human experiments, anti-CD34-FITC (0.5µL/well), anti-CD38-PerCP-Cy5.5 (0.1µL/well), anti-CD90-PE-Cy7 (0.25µL/well), and anti-CD45RA-PE (0.5µL/well) were used per well. After incubation, 100µL of PBS + 2%FCS was added to wells, and the plates were centrifuged 5 minutes at 4°C at 400*g* with low acceleration and medium deceleration. 100μL supernatant was aspirated and the cell pellet was resuspended in 200µL PBS+2%FCS + 0.1% PI + 0.25% Flow-Check Fluorspheres (Beckman Coulter). Samples were acquired in fast mode, and volumes of 100µL (for large colonies) or 150-170µL (for small colonies) were analyzed. Data were exported as FCS files and analyzed using FlowJo software. Cell number was corrected by bead count of Flow-Check (∼1000/µL). Megakaryocytes were identified as cells with high forward scatter and side scatter, as well as high CD150 and CD41 expression.

#### cDNA synthesis and quantitative RT-PCR

Approximately 7000-15000 fresh or cultured cells per condition were subjected to RNA extraction using an RNA-easy mini kit (QIAGEN). cDNA was synthesized using SuperScript VILO (Thermo Scientific Technology) in a final volume of 20µL, according to the manufacturer’s instruction.

qPCR was performed using SYBR Premix ExTaq^™^ IIa (TaKaRa Bio) according to manufacturer’s instruction. A mixture of 45μL SYBR Premix, 0.36μL each of forward and reverse primers, 1.8µL of Rox II dye, 41.48µL of distilled water, and 1µL of cDNA solution was established and 20µL was dispensed to each of 4 wells of an assay plate. PCR analysis was performed using an ABI 7500 Fast Real-Time PCR System (Applied Biosystems) under the following conditions: 95°C for 10s followed by 40 cycles of 95°C for 5s and 60°C for 34s. Expression levels were determined as 2^(Ct value – mean Ct value of GAPDH) and were normalized to control samples, unless otherwise stated.

#### cDNA microarray analysis

CD150^+^CD41^-^CD48^-^CD34^-^Flt3^-^LSK cells of pooled bone marrow from 10 (for 7-day cultures) or 20 (for 16-hour cultures) mice were sorted into SF-O3 medium and then and either lysed for the fresh sample or cultured. Cultured cells were centrifuged at 340 x *g* for 5 min at 4°C and lysed with 75µL RLT buffer + 0.75µL 2-ME. RNA extraction, cDNA synthesis, microarray analysis, and data normalization were outsourced to DNA Chip Research Inc. RNA was extracted using an RNeasy micro kit (QIAGEN). cDNA was synthesized and amplified from 2ng of extracted RNA using a WT-Ovation Pico RNA Amplification System (NuGen). Amplified cDNA was labelled using a Genomic DNA Enzymatic Labeling Kit (Agilent), followed by hybridization to the gene chip Mouse 8x60k (Agilent) using a Gene Expression Hybridization Kit (Agilent) and Gene Expression Wash Pack (Agilent).

### QUANTIFICATION AND STATISTICAL ANALYSIS

#### Statistical analysis

Data are presented as means ± SD, unless otherwise stated. For multiple comparisons, statistical significance was determined by Tukey’s multiple comparison test using the Tukey HSD function or one-way or two-way ANOVA using the anova and the aov function of R software. An unpaired two-tailed Student’s t test was used for experiments with two groups. In some transplantation assays with high variation, the Wilcoxon rank sum test was calculated using the wilcox.eact function of R software. False discovery rate (FDR) was calculated using the qvalue function of the qvalue package from Bioconductor (http://www.bioconductor.org).

#### cDNA microarray analysis

Scanning data and normalization were performed by DNA Chip Research Inc. The hybridized gene chip was scanned using DNA MicroArray Scanner (Agilent), and scanned images were analyzed with Feature Extraction Ver.9.5.3 (Agilent). Normalization was performed using GeneSpring software (Agilent). Raw data were first transformed to log_2_ scale with expression levels <1.0 set to 1.0. Gene expression levels across samples were normalized using the 75 percentile shift algorithm. Visualization of scatter plots of normalized expression with gene name annotation was performed using the ggplot2 package of R software. Genes with a log_2_ fold difference in expression of >1.5 were shown in dark blue, and genes of interest (i.e. fatty-acid-related genes) were in red. Genes with a “Not detected” flag were removed from the scatter plot.

#### Gene ontology analysis

Gene ontology (GO) analysis was performed using GeneSpring software (Agilent) for differentially expressed genes (DEG) between 2 samples. DEG was determined as genes with a fold difference of expression >2.0 between 2 samples.

#### GSEA

Normalized expression data were assessed using GSEA v2.0.13 software (Broad Institute). Gene sets were obtained from the Molecular Signatures Database v4.0 distributed at the GSEA website (http://www.broadinstitute.org/gsea/msigdb/index.jsp). The number of permutations was set to 1000. Gene sets with a nominal p-value <0.05 and a false discovery rate q-value <0.25 were considered statistically significant.

#### Ingenuity pathway analysis

Ingenuity pathway analysis (QIAGEN) was performed using microarray data from fresh HSCs in serum-free conditions (16 hours). The top 200 differentially-expressed genes were analyzed, and significantly enriched canonical pathways or upstream regulators in serum-free conditions were listed.

#### Estimation of doubling time

Doubling time in Figure S4S was estimated using following formula: 7 / log_2_ ((Mean of total cell count) / (Input cell number))

Upper estimate was:

7 / log_2_ ((Mean of total cell count -SD) / (Input cell number))

Lower estimate was:

7 / log_2_ ((Mean of total cell count +SD) / (Input cell number))

When (Mean-SD)/Input <1, the result was denoted as NA.

#### Single cell qPCR analysis

Data were analyzed using Biomark qPCR analysis software (Fluidigm). Melting curves showing low quality (Quality threshold <0.65) and a Peak Ratio Threshold of <0.8 were interpreted as not detected. The baseline was corrected by linear correction. Data were exported to a .csv file and downstream analysis was performed using R software. Cells lacking expression of Gapdh, Hprt1 or Actb were excluded, leaving 46 of 48 fresh cultures and 42 of 48 cultured HSCs. Genes not expressed in any cell (such as Cdkn2a, Cdkn2b, Csf1r, and Il7ra) were also excluded from analysis, leaving 92 of 96 genes. For each sample, Ct values were normalized to expression levels of housekeeping genes by subtracting average Ct values of Actb and Gapdh (normalized Ct values were designated ΔCt values). ΔCt values for not detected genes were set to the maximum ΔCt value of each gene + 3.5. For principal component analysis, the prcomp function implemented in R software was used on ΔCt values. Housekeeping genes (Actb, Gapdh, and Hprt1) were removed prior to analysis. Data scaling was not applied. The first 2 principal components were used to generate the plot. Hierarchical clustering with Euclidean distance and complete linkage clustering was performed on the correlation coefficient matrix of selected genes calculated by the cor function with Pearson’s correlation coefficient. Genes with a p-value <0.05 (calculated from ΔCt values using the t.test function) between fresh and cultured HSCs were selected for the cluster analysis. Hierarchical clustering was performed using the hclust package and the heatmap.2 function from the gplots package. All figures, including violin plots of ΔCt values, PCA plots, and hierarchical clustering, were visualized using the ggplot2 package. For visualization of expression levels in violin plots, the maximum ΔCt of each gene + 3.5 – ΔCt was shown, setting expression levels of not detected genes to zero. The false-discovery rate of differences between expression levels in fresh versus cultured cells was calculated using the qvalue function in the qvalue package after creating a list of p-values for each gene using the t.test function.

#### Generation of 2D and 3D plots for cytokine response

Cell number or fold-change of phosphoproteins under various cytokine conditions was visualized followed by smoothing using R software, so that the global landscape of HSC behavior in the cytokine concentration space is readily recognized. Local regression was performed to fit a polynominal surface to data points using the loess function with parameters set as, degree = 2, and span = 0.25. X-, and Y-axes were expanded from 7 x 7 matrix to a 61 x 61 matrix, and data prediction was performed using the predict function. When data were log scale (as in total cell number or megakaryocyte number), the raw data value was increased by 1 to bring the data point above 0. 2D contour plots were generated using the ggplot2 package, and 3D plots were generated using the rgl package.

### DATA AND SOFTWARE AVAILABILITY

cDNA microarray data generated here are accessible in the GEO database under the accession number GSE117515 and GSE117516. All software packages and methods used in this study have been properly detailed and referenced under “QUANTIFICATION AND STATISTICAL ANALYSIS”.

## SUPPLEMENTAL FIGURE LEGENDS

**Figure S1, related to Figure 1.**
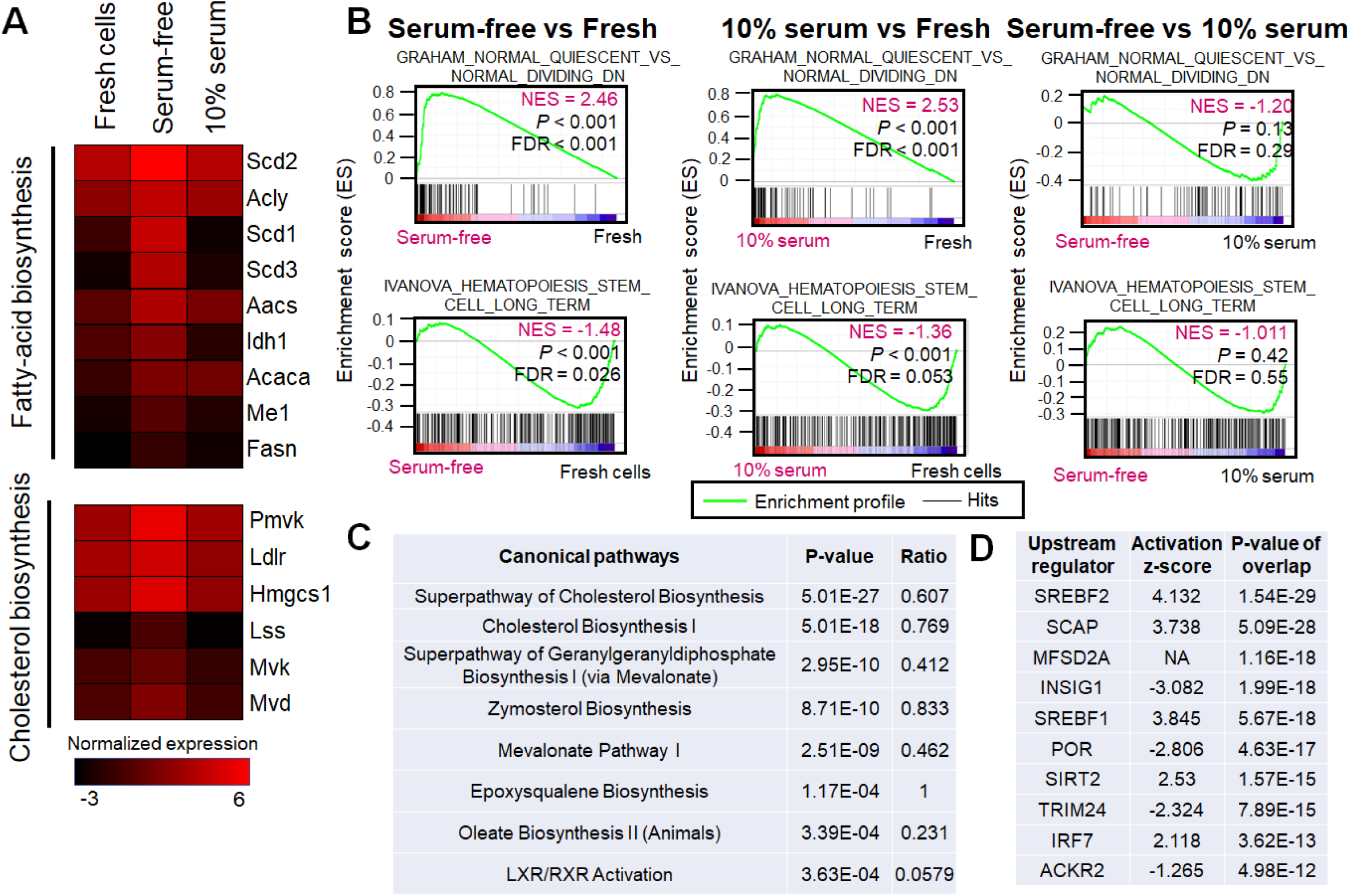
(**A**) Heat map showing expression levels of fatty acid and cholesterol biosynthesis genes based on cDNA microarray analysis. Freshly isolated CD150^+^CD48^-^CD41^-^Flt3^-^CD34^-^ LSK cells (Fresh HSC), HSCs cultured 16 hours with (10% serum) or without (Serum-free) serum were subjected to cDNA microarray analysis in Figure 1A. Cytokines (SCF and TPO at 100ng/mL each) were supplemented in each condition. (**B**) Gene set enrichment analysis (GSEA) of cDNA microarray data of fresh HSCs and HSCs cultured 16 hours with or without 10% serum. FDR, false discovery rate. NES, normalized enrichment score. (**C**) Canonical pathways over-represented in serum-free culture based on Ingenuity Pathway Analysis (IPA) using cDNA microarray data in Figure 1A. (**D**) Upstream regulators over-represented in serum-free culture based on Ingenuity Pathway Analysis (IPA) using cDNA microarray data.

**Figure S2, related to Figure 2.**
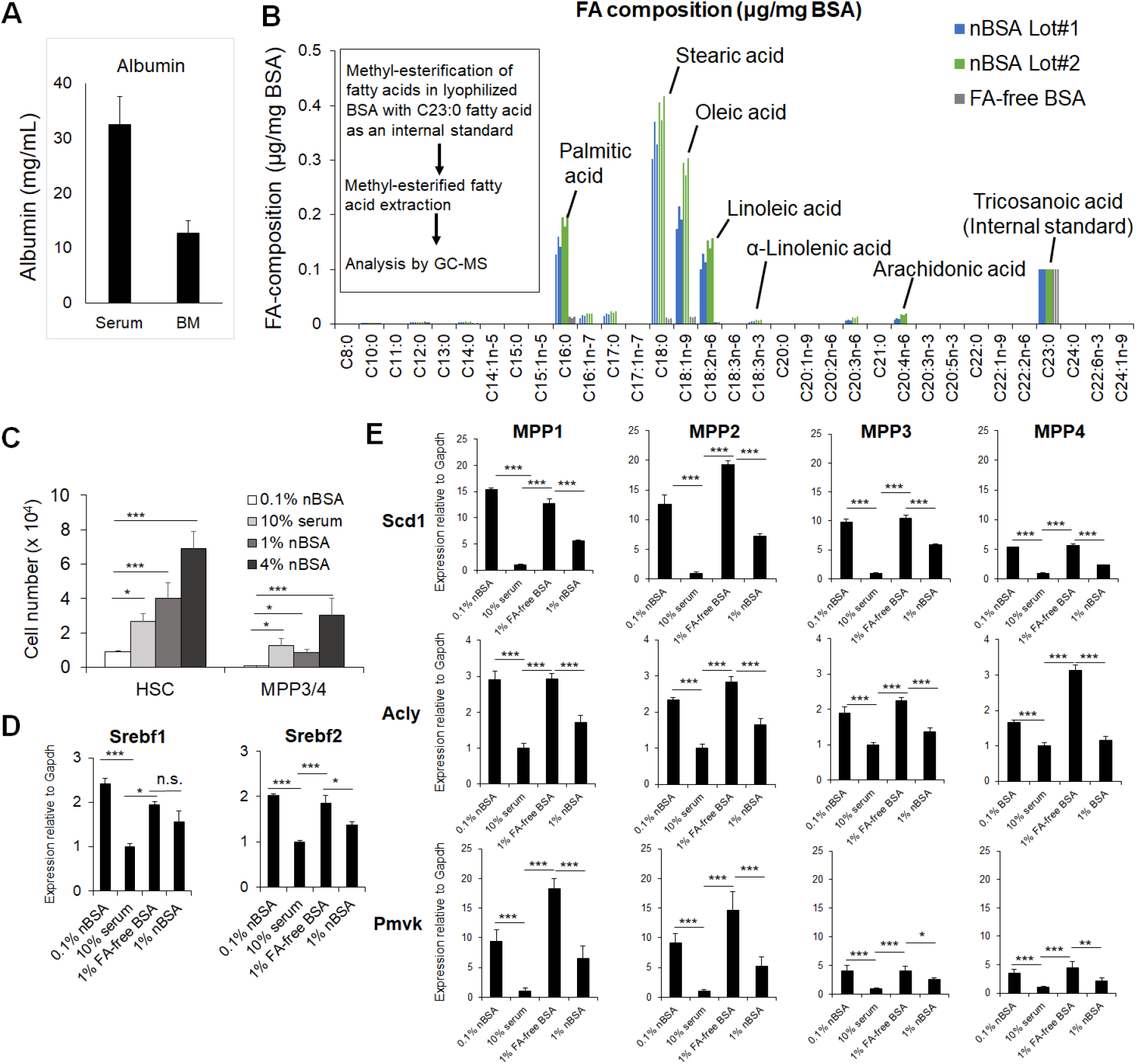
(**A**) Albumin concentration of murine serum and bone marrow extracellular fluid measured by ELISA (mean ± SD, n =3 from 3 mice). (**B**) Lipidomic identification of BSA binding to fatty acid. Experimental design (left) and FA composition of crystallized native BSA (Lots #1 and #2), and FA-free BSA, as measured by gas chromatography/mass spectrometry (GC-MS) (right, n =3 per each condition). (**C**) After culturing either 100 CD150^+^CD48^-^CD41^-^Flt3^-^CD34^-^ LSK cells (HSCs) or CD150^-^CD48/CD41^+^ LSK cells (MPP3/4) for 7 days, we determined the total number of cells in indicated conditions in SF-O3 with SCF and TPO at 100ng/mL each (mean ± SD, n = 4). \**P* < 0.05, \*\**P* < 0.01, and \*\*\**P* < 0.001. n.s., not significant among each of HSC or MPP by Tukey-Kramer multiple comparison test. (**D**) qPCR analysis of Srebf1 and Srebf2 mRNAs in CD150^+^CD41^-^CD48^-^Flt3^-^CD34^-^ LSK cells cultured for 16 hours. Indicated cultures were supplemented with SCF and TPO at 100ng/mL each. nBSA, native bovine serum albumin; FA-free BSA, 1% w/v fatty-acid-free BSA. Values were normalized to Gapdh expression and expressed as fold-induction compared to levels detected in 10% serum (mean ± SD, n = 4 technical replicate). \**P* < 0.05, and \*\*\**P* < 0.001. n.s., not significant by Tukey-Kramer multiple comparison test. (**E**) qPCR analysis of fatty acid- or cholesterol-related mRNAs of MPP fractions cultured 16 hours. Cultures contained SCF and TPO at 100ng/mL each. nBSA, native bovine serum albumin. FA-free BSA: 1% w/v fatty-acid-free BSA. Values were normalized as above and expressed as fold-induction compared to levels detected in 10% serum (mean ± SD, n = 4 technical replicate). \**P* < 0.05, \*\**P* < 0.01, and \*\*\**P* < 0.001. n.s., not significant by Tukey-Kramer multiple comparison test. MPP1, CD150^-^CD41^-^CD48^-^Flt3^-^ LSK cells. MPP2, CD150^+^CD41/CD48^+^Flt3^-^ LSK cells. MPP3, CD150^-^CD41/CD48^+^Flt3^-^ LSK cells. MPP4, Flt3^+^ LSK cells.

**Figure S3, related to Figure 3.**
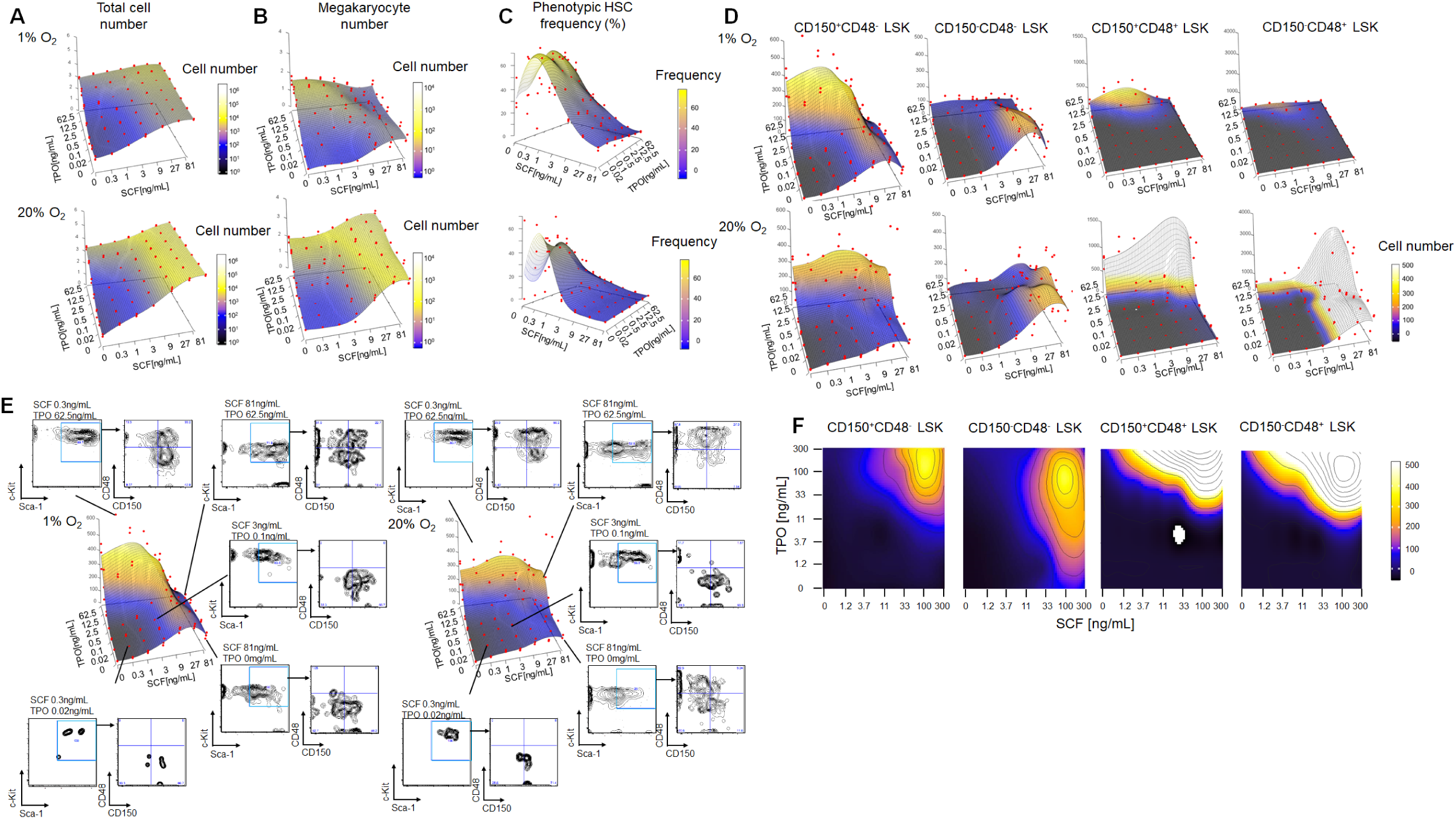
(**A-D**) 3D plots with data points (red) used to construct the predicted planes shown in Figures 3A-3D. (**E**) Representative flow cytometric plots for CD150^+^CD48^-^ LSK culture shown in Figure S3D. (**F**) CD150^+^CD41^-^CD48^-^Flt3^-^CD34^-^ LSK cells (100/well) were cultured 7 days in 0.1% BSA and O_2_ conditions with 49 (7 x 7) different SCF and TPO concentrations (ranging from 0 to 300ng/mL). A MACSQuant analyzer was used to determine indicated cell numbers. Data were smoothened, reshaped to a 61 x 61 matrix, and presented as contour plots with a color key (n = 3 from 1 experiment)

**Figure S4, related to Figure 4.**
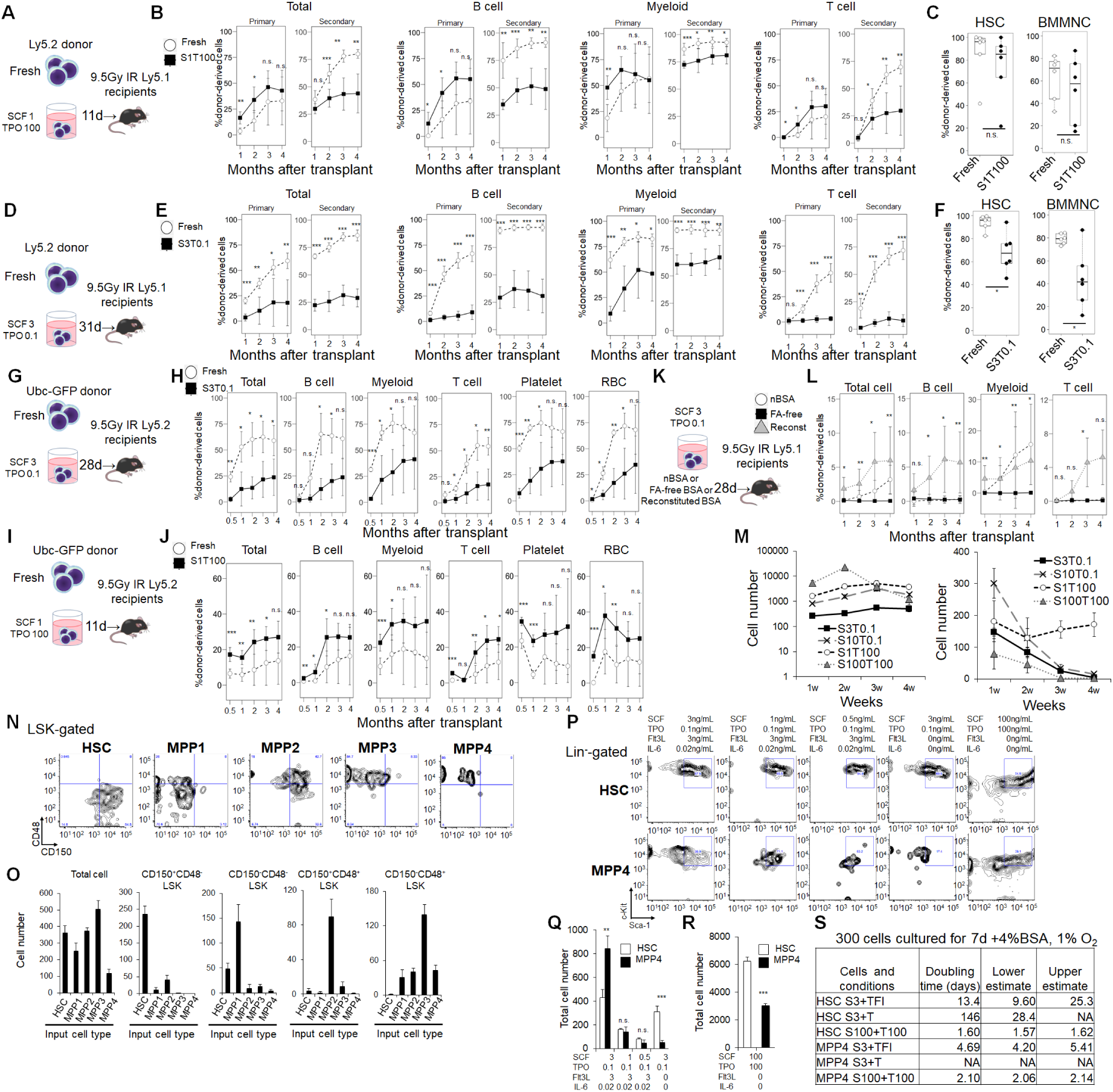
(**A-L**) BMT assay for fresh and/or cultured HSCs in five different conditions. (**A**) Experimental design. CD150^+^CD41^-^CD48^-^Flt3^-^CD34^-^ LSK cells (HSCs) from Ly5.2^+^ mice were cultured 11 days. For primary BMT, 100 Freshly-isolated HSCs (Fresh), or 100 HSCs cultured with 1ng/mL SCF plus 100ng/mL TPO (S1T100) in 1% O_2_ were transplanted into lethally-irradiated (9.5Gy) Ly5.1^+^ recipients together with 4 x 10^5^ Ly5.1^+^ bone marrow mononuclear cells, and donor-derived chimerism of peripheral blood was examined. For secondary BMT, 2 x 10^6^ bone marrow mononuclear cells were transplanted into recipients after assessment for chimerism in recipients’ bone marrow. (**B**) Peripheral blood chimerism of HSCs cultured 11 days in (**A**) (mean ± SD, n = 6 per each group). \**P* < 0.05, \*\**P* < 0.01, and \*\*\**P* < 0.001. n.s., not significant by two-tailed Student’s t test. (**C**) Bone marrow chimerism of donor-derived cells 4 months after primary BMT of (**B**) (n = 6 per group). \**P* < 0.05, \*\**P* < 0.01, and \*\*\**P* < 0.001. n.s., not significant by two-tailed Student’s t test. (**D**) Experimental design. Replicate of Figure 4G with a different BSA lot and a longer culture period. CD150^+^CD41^-^CD48^-^Flt3^-^CD34^-^ LSK cells (HSC) from Ly5.2^+^ mice were cultured 31 days. For primary BMT, 500 Freshly-isolated HSCs (Fresh) or 500 HSCs cultured with 3ng/mL SCF plus 100ng/mL TPO (S3T0.1) at 1% O_2_ were transplanted into lethally-irradiated (9.5Gy) Ly5.1^+^ recipients together with 4 x 10^5^ Ly5.1^+^ bone marrow mononuclear cells, and donor-derived chimerism of peripheral blood was examined. For secondary BMT, 2 x 10^6^ bone marrow mononuclear cells were transplanted into recipients as described after assessment of chimerism in primary recipients’ bone marrow. Conditions with SCF and TPO at 100ng/mL each were not tested. (**E**) Peripheral blood chimerism of HSCs cultured 31 days in (**D**) (mean ± SD, n = 6 per each group). \**P* < 0.05, \*\**P* < 0.01, and \*\*\**P* < 0.001. n.s., not significant by two-tailed Student’s t test. (**F**) Bone marrow chimerism of donor-derived cells 4 months after primary BMT of (**E**) (n = 6 per group). \**P* < 0.05, \*\**P* < 0.01, and \*\*\**P* < 0.001. n.s., not significant by two-tailed Student’s t test. (**G**) Experimental design. CD150^+^CD41^-^CD48^-^Flt3^-^ LSK cells (HSCs) from Ubc-GFP mice were cultured 28 days and then BMT was performed as in (**D**). (**H**) Peripheral blood chimerism of HSCs cultured 28 days in (**G**) (mean ± SD, n = 3 for Fresh and n = 4 for S3T0.1). \**P* < 0.05, \*\**P* < 0.01, and \*\*\**P* < 0.001. n.s., not significant by two-tailed Student’s t test. (**I**) Experimental design. CD150^+^CD41^-^CD48^-^Flt3^-^ LSK cells (HSC) from Ubc-GFP mice were cultured 11 days and then BMT was performed as in (**A**). (**J**) Peripheral blood chimerism of HSCs cultured 28 days (**I**) (mean ± SD, n = 6 per group). (**K**) Experimental design. CD150^+^CD41^-^CD48^-^Flt3^-^CD34^-^ LSK cells (HSCs) from Ly5.2^+^ mice were cultured 28 days. An equivalent of 500 HSCs cultured with SF-O3 plus 3ng/mL SCF and 100ng/mL TPO (S3T0.1) in 1% O_2_ under indicated conditions were transplanted into lethally-irradiated (9.5Gy) Ly5.1^+^ recipients together with 4 x 10^5^ Ly5.1^+^ bone marrow mononuclear cells, and donor-derived chimerism of peripheral blood was examined. Reconst. BSA: 4% w/v fatty acid-free BSA reconstituted with sodium palmitate and sodium oleate at 200μg/mL each. (**L**) Peripheral blood chimerism of HSCs cultured for 28 days in (**K**) (mean ± SD, n = 6 for nBSA, n= 6 for FA-free, and n = 5-6 for Reconst.). \**P* < 0.05, \*\**P* < 0.01, and \*\*\**P* < 0.001. n.s., not significant between nBSA and Reconst. BSA. Wilcoxon’s rank sum test was used rather than a t test due to high variation. (**M**) Total cell number (left) and the number of CD150^+^CD48^-^ LSK cells (right) after 1, 2, 3, and 4 weeks of culture of 300 CD150^+^CD41^-^CD48^-^Flt3^-^CD34^-^ LSK cells in indicated conditions (mean ± SD, n = 4 for each group) (**N and O**) Indicated HSPC fractions were cultured in 1% O_2_ for 5 days in 3ng/mL SCF and 0.1ng/mL TPO plus 4% nBSA and then analyzed by flow cytometry. 300 input cells were used. HSCs: CD150^+^CD41^-^ CD48^-^Flt3^-^CD34^-^ LSK cells. MPP1: CD150^-^CD41^-^CD48^-^Flt3^-^ LSK cells. MPP2: CD150^+^CD41/CD48^+^Flt3^-^ LSK cells. MPP3: CD150^-^CD41/CD48^+^Flt3^-^ LSK cells. MPP4: Flt3^+^ LSK cells. (**N**) Representative plots for flow cytometry. (**O**) Number of cells with indicated surface marker profile (mean ± SD, n = 4). (**P-S**) CD150^+^CD41^-^CD48^-^Flt3^-^CD34^-^ LSK cells (HSCs) and Flt3^+^ LSK cells (MPP4) were cultured 7 days in indicated cytokine conditions in 1% O_2_. 100 input cells were used. Medium contained 4% nBSA. Shown are representative plots for flow cytometry (**P**), and total cell number in low (**Q**) or high (**R**) cytokine conditions (mean ± SD, n = 4). \**P* < 0.05, \*\**P* < 0.01, and \*\*\**P* < 0.001. n.s., not significant by two-tailed Student’s t test between HSC and MPP4. (**S**) Doubling time of HSCs and MPP4s, as estimated from observations shown in (**Q**), and (**R**).

**Figure S5, related to Figure 5.**
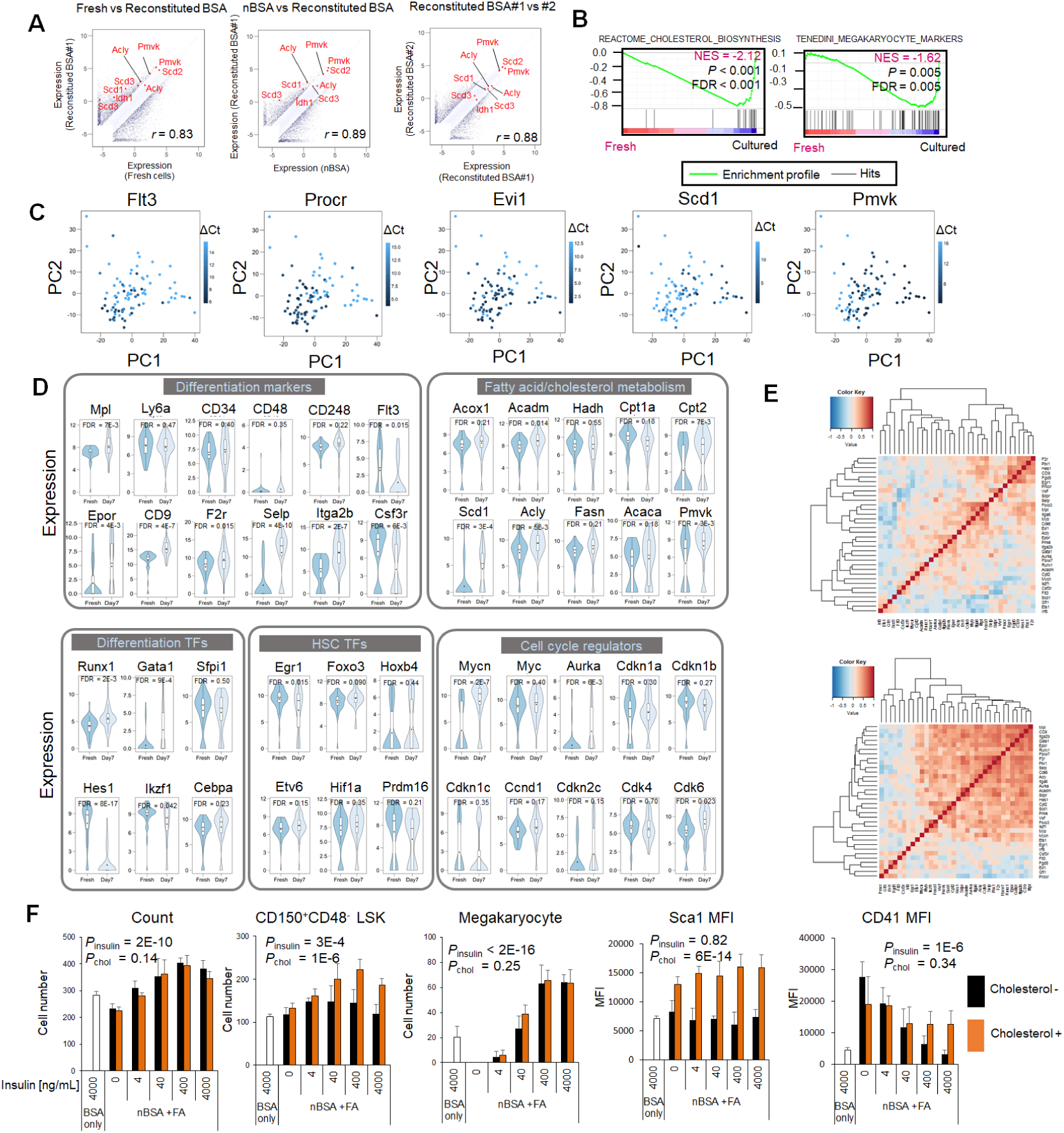
(**A**) Gene expression levels in indicated samples from microarray data shown in Figure 5A. Differentially expressed genes are in dark blue, and fatty acid-synthesis-related genes are in red. Correlation coefficient *r* is noted. (**B**) Gene set enrichment analysis (GSEA) of cDNA microarray data of fresh HSCs and HSCs cultured 7 days (a mixture of nBSA and reconstituted BSA samples) in Figure 5A. FDR, false discovery rate. NES, normalized enrichment score. (**C**) Principal component analysis of the sc-qPCR data in Figure 5D, overlaid by color keys indicating ΔCt values for each gene. (**D**) Violin plots of gene expression levels of HSC- and differentiation-related genes as measured by sc-qPCR in Figure 5D. False discovery rate (FDR) was calculated after analysis using a two-tailed Student’s t test. (**E**) Correlation coefficient matrix with hierarchical clustering of differentially expressed genes between fresh and cultured HSCs. sc-qPCR data of fresh (top) and cultured (bottom) HSCs were separately analyzed. (**F**) Effects of insulin and cholesterol on HSC phenotypes. CD150^+^CD48^-^CD41^-^ LSK cells were cultured 7 days in 4% nBSA (BSA only) or 4% nBSA plus a fatty-acid mixture (palmitate, oleate, linoleate, and stearate at a 4:3:2:1 ratio), with or without cholesterol in 1% O_2_. Insulin was added at indicated concentrations (mean±SD, n=4). The number of total cells, CD150^+^CD48^-^ LSK cells, and megakaryocytes, and mean fluorescence intensity (MFI) of Sca-1, and CD41 were measured. P values were calculated by two-way ANOVA to examine effects of insulin (*P*_ins_) and cholesterol (*P*_cho_). BSA only plus 4000ng/mL insulin samples were removed from the ANOVA.

**Figure S6, related to Figure 6.**
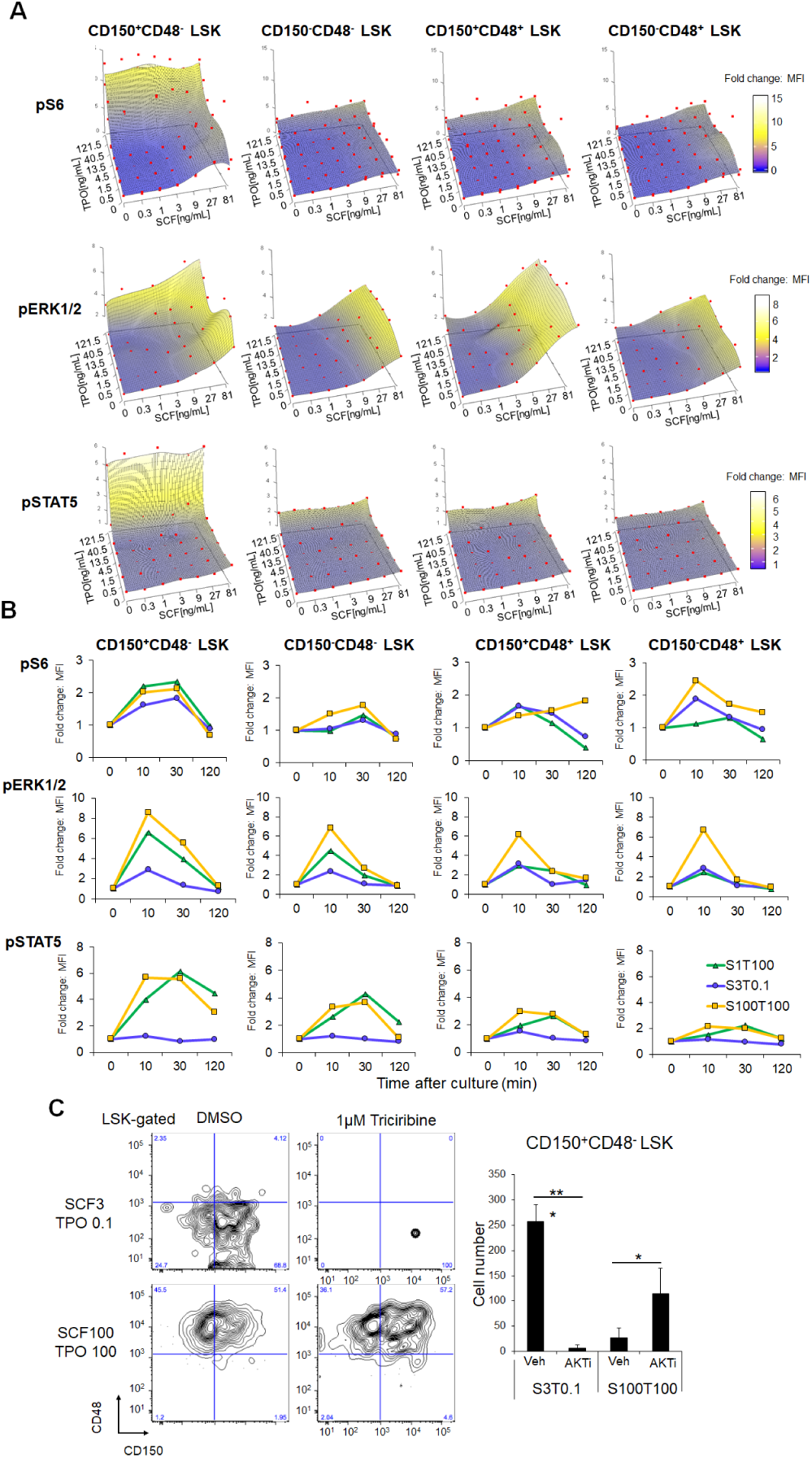
(**A**) 3D plots with data points (red) used to predict the plane shown in Figure 6A. (**B**) Phosphorylation status of S6, ERK1/2, and STAT5 in hematpoietic stem/progenitors based on intracellular flow cytometry following cytokine exposure. Bone marrow cells were incubated for 0 (untreated control), 10, 30, and 120 minutes with indicated cytokine conditions. Shown is fold-change of mean fluorescent intensity (MFI) relative to untreated control of phosphoproteins in indicated fractions (n = 1 for each group). (**C**) Representative flow cytometric plots (left) and the number of CD150^+^CD48^-^ LSK cells after 8 days of culture with 1μM of the AKT inhibitor Triciribine (right) (mean ± SD, n=4). \**P* < 0.05, \*\**P* < 0.01, \*\*\**P* < 0.001 by Tukey-Kramer multiple comparison test.

**Figure S7, related to Figure 7.**
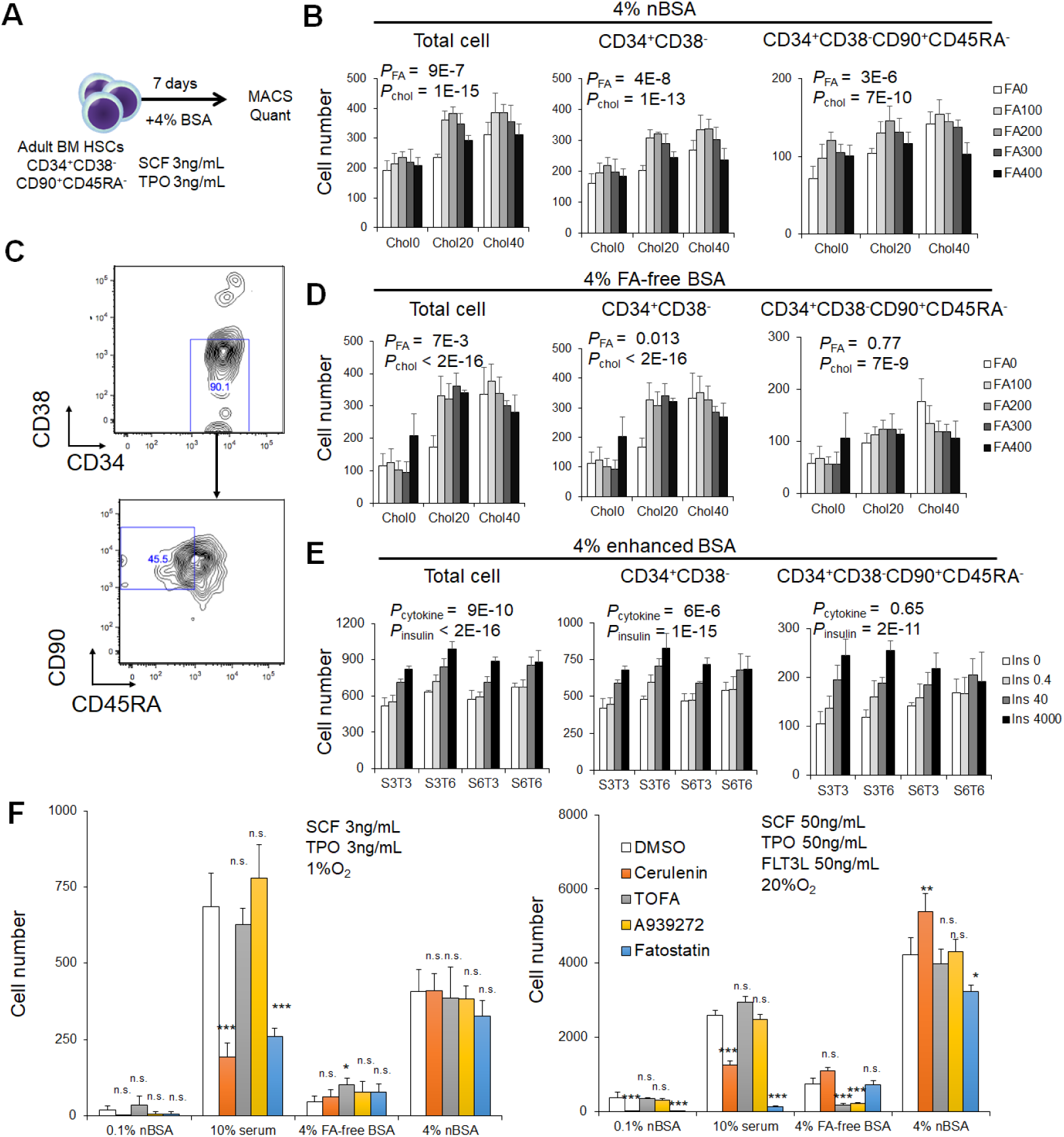
(**A-E**) Frozen adult bone marrow CD34^+^ cells were thawed and stained for surface markers, and CD34^+^CD38^-^CD90^+^CD45RA^-^ cells were sorted and a total of 600 cells were cultured 7 days in indicated conditions. Experimental design (**A**). To assess effects of fatty acid and cholesterol, human HSCs were cultured in indicated fatty acid (μg/mL) and cholesterol (μg/mL) concentrations in 1% O2. Medium was SF-O3+4% nBSA plus a FA mixture (palmitate, oleate, linoleate, and stearate at a 4:3:2:1 ratio). Media contained SCF and TPO at 3ng/mL each. The number of total cells and CD34^+^CD38^-^ and CD34^+^CD38^-^ CD90^+^CD45RA^-^ cells was determined (mean ± SD, n=4). P values were calculated by two-way ANOVA to examine the influence of fatty acid (*P*_FA_) and cholesterol (*P*_chol_) (**B**). Representative plots showing flow cytometry analysis of HSCs cultured in 200μg/mL FA, 20μg/mL cholesterol, and SCF and TPO at 3ng/mL in 1% O_2_ (**C**). HSCs were cultured in indicated fatty acid (μg/mL), and cholesterol (μg/mL) concentrations in 1% O_2._ Medium was SF-O3 plus 4% FA-free BSA plus a FA mixture (palmitate, oleate, linoleate, and stearate at a 4:3:2:1 ratio). SCF and TPO were at 3ng/mL each. Total, CD34^+^CD38^-^, and CD34^+^CD38^-^ CD90^+^CD45RA^-^ cells were measured (mean ± SD, n=4). P values were calculated by two-way ANOVA to examine the influence of fatty acid (*P*_FA_) and cholesterol (*P*_chol_) (**D**). To assess insulin dependency, 1000 human CD34^+^CD38^-^CD90^+^CD45RA^-^ cells were cultured 7 days in indicated insulin concentrations. Culture medium was αMEM plus 4% enhanced BSA (palmitate, oleate, linoleate, and stearate at a 4:3:2:1 ratio and 20μg/mL cholesterol). S3: SCF 3ng/mL. S6: SCF 6ng/mL, T3: TPO 3ng/mL. T6: TPO 6ng/mL. The number of total, CD34^+^CD38^-^, and CD34^+^CD38^-^CD90^+^CD45RA^-^ cells was measured (mean ± SD, n=4). P values were calculated by two-way ANOVA to examine the influence of cytokine (*P*_cytokine_) and insulin (*P*_insulin_) (**E**). (**F**) HSC proliferation or survival in the presence of fatty acid synthesis inhibitors, and in the presence or absence of fatty acids. A total of 600 CD34^+^CD38^-^CD90^+^CD45RA^-^ cells were cultured 7 days with indicated inhibitors. Medium was SF-O3 supplemented with either 0.1% nBSA, 10% serum, 4% FA-free BSA, or 4% nBSA. Also added was SCF and TPO at 3ng/mL each (left panel) or SCF, TPO and FLT3L each at 50ng/mL (right panel). The lower cytokine group was cultured in 1% O_2_, and the higher cytokine group was cultured in 20% O_2_ conditions.

## KEY RESOURCES TABLE

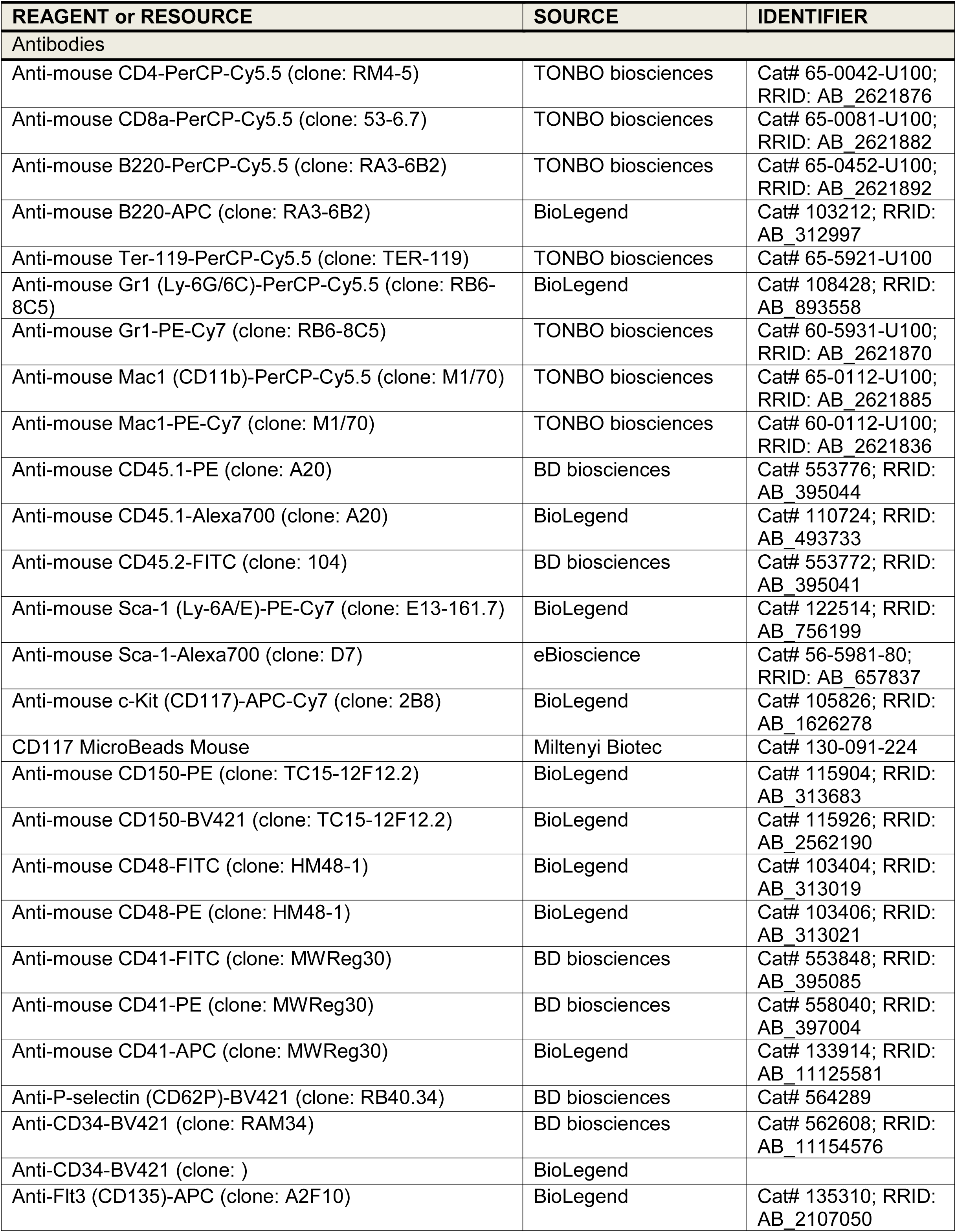
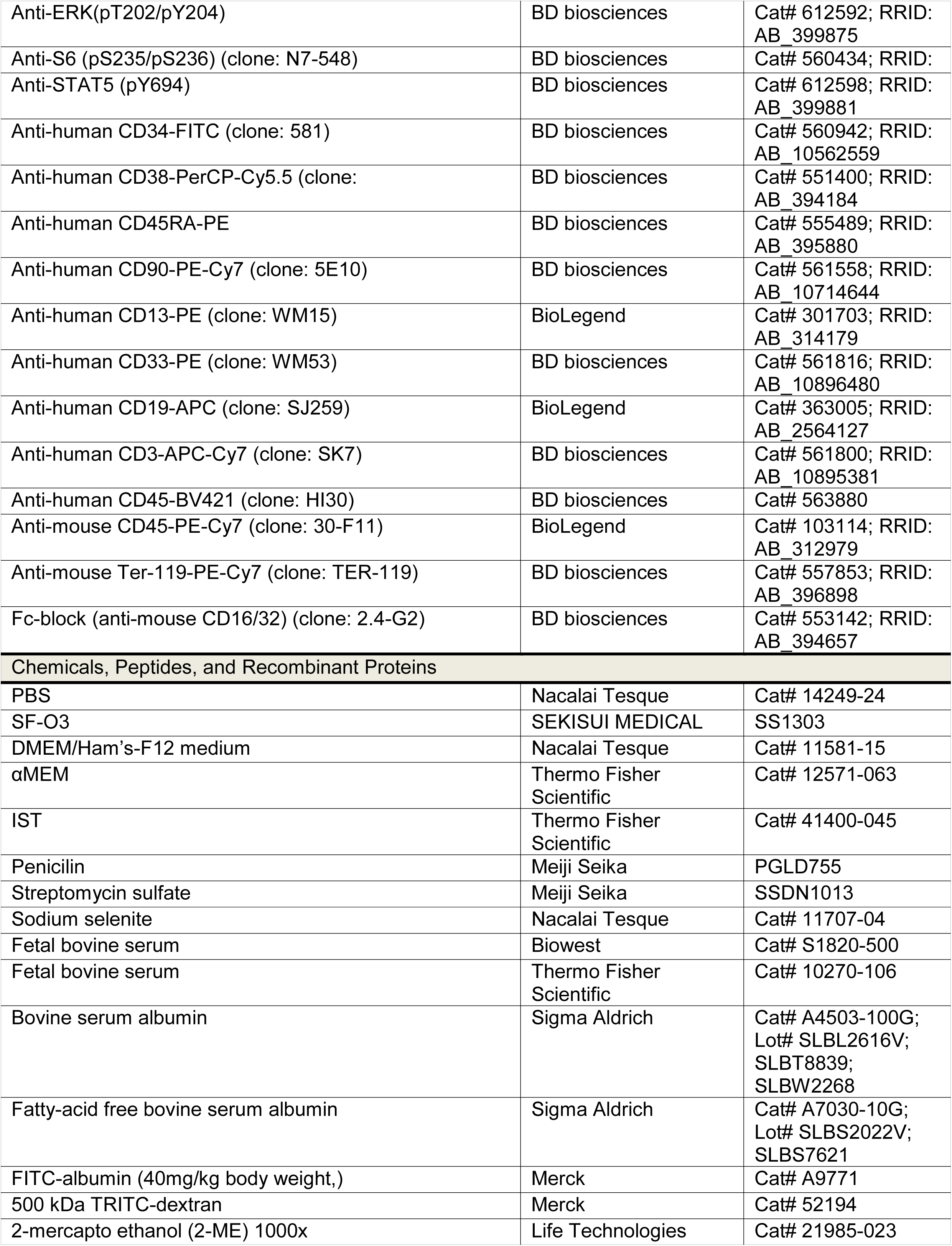
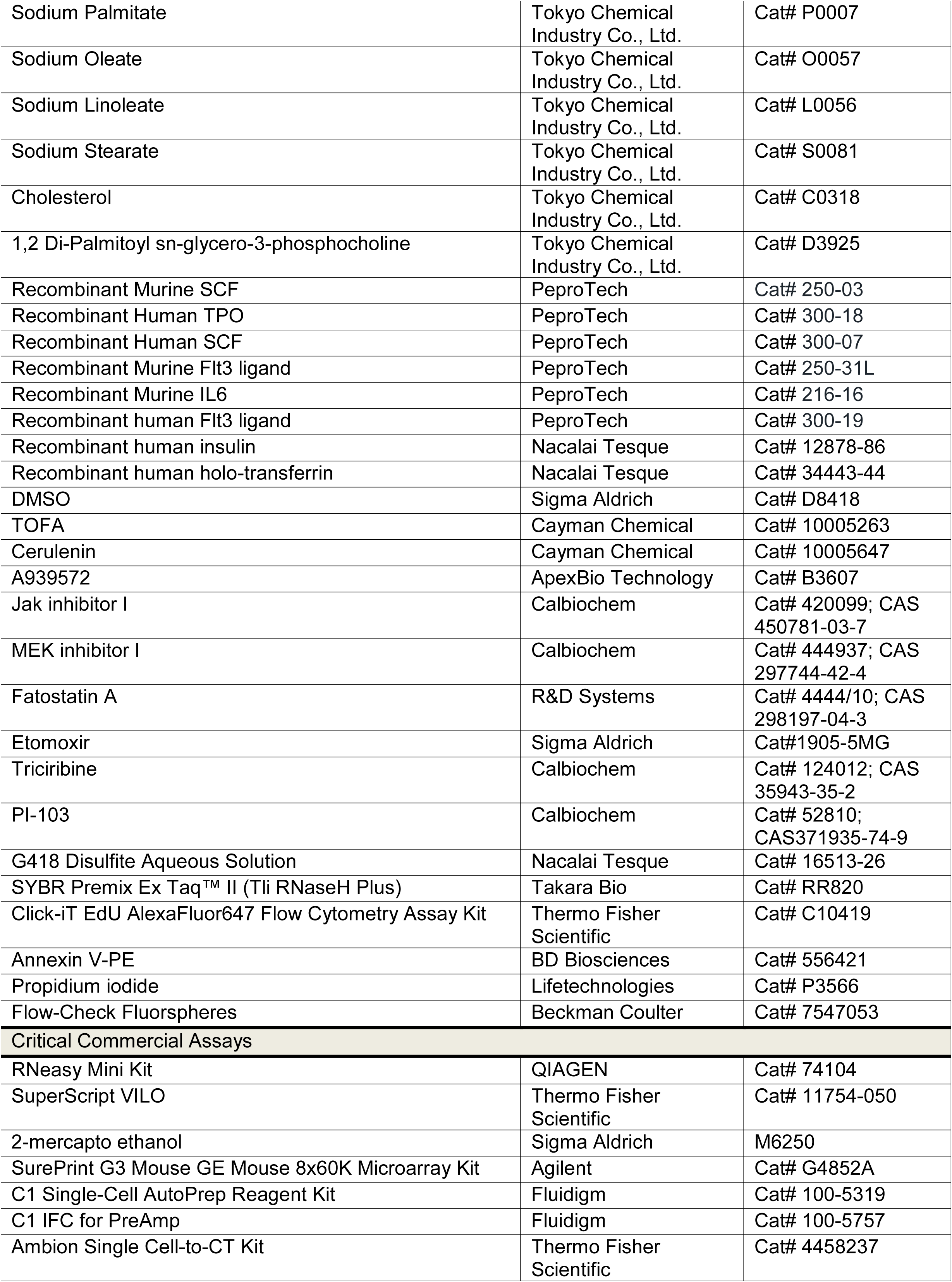
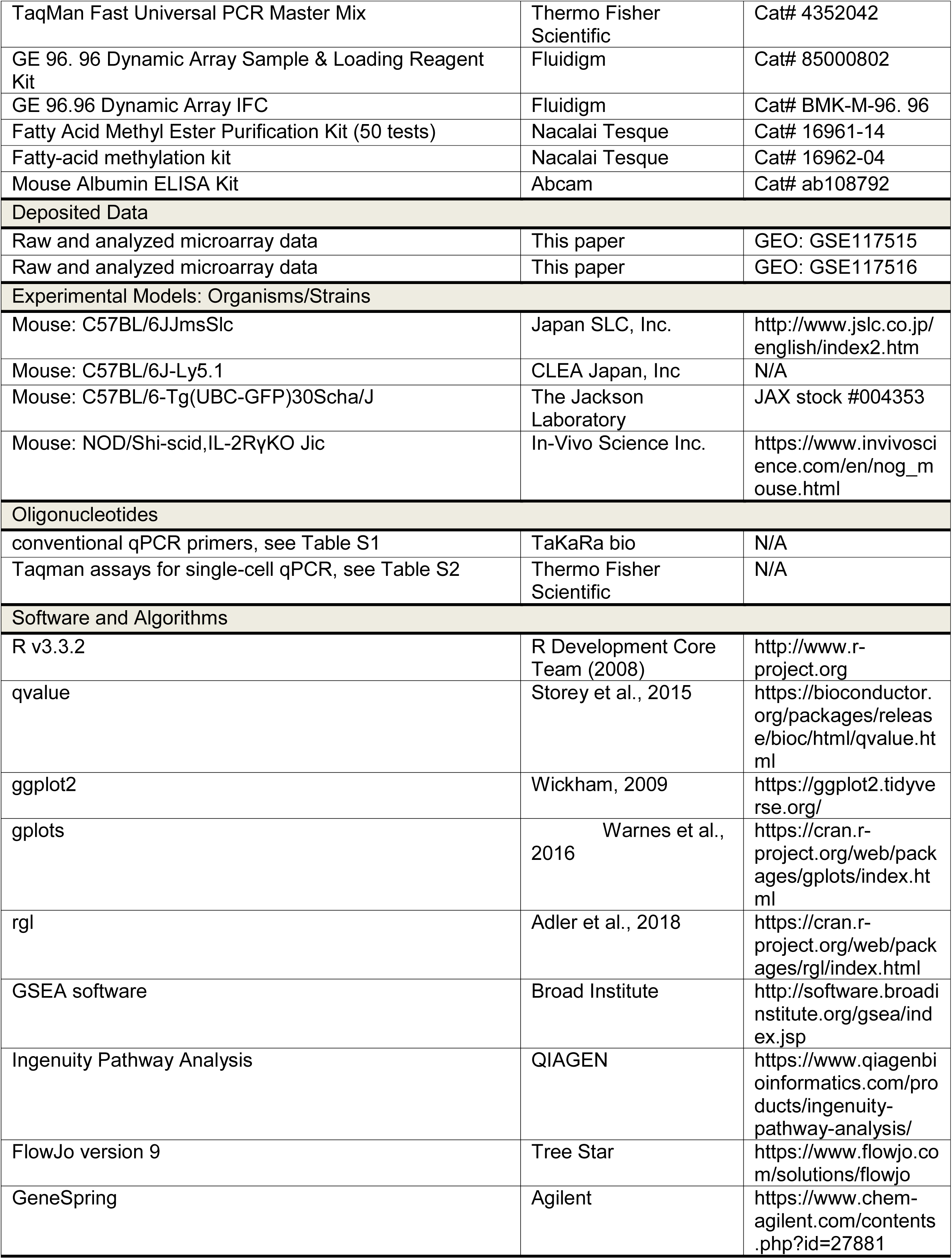

